# Characterization of neurogenic niches in the telencephalon of juvenile and adult sharks

**DOI:** 10.1101/730721

**Authors:** A Docampo-Seara, S Pereira-Guldrís, N Sánchez-Farías, S Mazan, MA Rodríguez, E Candal

## Abstract

Neurogenesis is a multistep process by which progenitor cells become terminally differentiated neurons. Adult neurogenesis has gathered increasing interest with the aim of developing new cell-based treatments for neurodegenerative diseases in humans. Active sites of adult neurogenesis exist from fish to mammals, although in the adult mammalian brain the number and extension of neurogenic areas is considerably reduced in comparison to non-mammalian vertebrates, and they become mostly reduced to the telencephalon. Much of our understanding in this field is based in studies on mammals and zebrafish, a modern bony fish. The use of the cartilaginous fish *Scyliorhinus canicula* (representative of basal gnathostomes) as a model expands the comparative framework to a species that shows highly neurogenic activity in the adult brain. In this work, we studied the proliferation pattern in the telencephalon of juvenile and adult specimens of *S. canicula* by using antibodies against the proliferation marker PCNA. We have characterized proliferating niches by using stem cell markers (Sox2), glial markers (GFAP, BLBP and GS), intermediate progenitor cell markers (Dlx2 and Tbr2) and markers for migrating neuroblasts (DCX). Based in the expression pattern of these markers, we demonstrate the existence of different cell subtypes within the PCNA immunoreactive zones including non-glial stem cells, glial progenitors, intermediate progenitor-like cells and migratory neuroblasts, which were widely distributed in the ventricular zone of the pallium, suggesting that the main progenitor types that constitute the neurogenic niche in mammals are already present in cartilaginous fishes.

## Introduction

Neurogenesis can be defined as a series of developmental events leading to a new neuron (Götz and Huttner, 2005; Espósito et al. 2005; Hevner et al. 2006). This definition involves the existence of progenitor cells that progressively restrict their proliferative potential and number of possible fates to become a terminally differentiated neuron (Hevner et al. 2006). Despite it was believed that the neurogenic process was only restricted to the developmental period, in the past 60 years it has been discovered that, in some species, adult individuals also present constitutive neurogenesis. In the mammalian brain, adult neurogenesis in the telencephalon takes place mainly in two regions: the subgranular zone (SGZ) of the dentate gyrus of the hippocampus (first described by Altman, 1963) and the subventricular zone (SVZ) of the lateral walls of the telencephalic ventricles (first described by Altman, 1969). Despite adult neurogenesis in the rodent brain has been largely proved, in primates this issue remains, to date, controversial (Boldrini et al. 2018; Sorrells et al. 2018; reviewed by Kempermann et al. 2018).

In the adult telencephalic niche of mammals, new-born neurons are generated from adult neural stem cells (aNSCs) also termed adult neural progenitor cells (Ming and Song, 2011; Martínez-Cerdeño and Noctor, 2018). Adult progenitors in mouse have been extensively characterized. Two models about the identity of these precursors were initially postulated (Ming and Song, 2011): (1) progenitor cells are radial glial cells that express both astroglial and stem cell markers (Álvarez-Buylla and Lim, 2004; Ma et al. 2009); (2) progenitor cells are not radial glial cells and they express stem cell markers such as *Sex determining region Y-box 2* (Sox2; Suh et al. 2007). Later, it has been discovered that both models are not mutually excluding but rather complementary, revealing the wide diversity of adult progenitor types (Bonaguidi, et al. 2012), and the need to deepen in the characterization of progenitor cells in the adult brain.

Nowadays, it is accepted that adult progenitor cells in the telencephalon of mammals can be subdivided in radial glia-like and non-radial progenitors. Radial glia-like progenitor cells have the capacity of self-renewal, show long-term maintenance of the undifferentiated state and generate different kind of neurons (Bonaguidi et al. 2016). Radial glia-like progenitor cells occasionally divide and generate non-radial progenitors (Bonaguidi et al. 2012). However, they normally exhibit a relatively quiescent state. This type of cells is known as B cells in the SVZ and as Type-1 cells in the SGZ (Doetsch et al. 1997, 1999; Seri et al. 2004; Ming and Song, 2011; Bond et al. 2015; Bonaguidi et al. 2016; Lim and Álvarez-Buylla, 2016). These progenitors express the glial fibrillary acidic protein (GFAP), the brain lipid binding protein (BLBP), glutamine synthase (GS) and Sox2, among others (Götz, 2013). On the other hand, non-radial progenitor cells are transit amplifying cells (Martínez-Cerdeño and Noctor, 2018) or intermediate progenitor cells (IPCs). IPCs are actively dividing cells that lack radial processes and they express proliferating and neuronal lineage markers that depend on their future phenotype (Suh et al. 2007; Lugert et al. 2010): GABAergic progenitors express the *Distal-less* homolog homeobox gene *Dlx2* and glutamatergic progenitors express the T-box transcription factor *Tbr2* (Hodge et al. 2012; Lim and Álvarez-Buylla, 2016). These cells are known as C cells in the SVZ and as Type-2 cells in the SGZ (Doetsch et al. 1997, 1999; Seri et al. 2004; Steiner et al. 2006; Ming and Song, 2011; Bond et al. 2015; Bonaguidi et al. 2016; Lim and Álvarez-Buylla, 2016). IPCs undergo mitosis generating more IPCs or two migratory neuroblasts. These neuroblasts leave the neurogenic niche and migrate to their final destinations in the brain. In the case of the SVZ, these neuroblasts are called A cells and migrate following a particular tangential pathway to the olfactory bulb called rostral migratory stream (RMS; reviewed by Lim and Álvarez-Buylla, 2016). In the SGZ, these cells are referred to as Type-3 cells and migrate locally to their final destination on the hippocampus. Both neuroblasts of the SVZ and SGZ express the same lineage markers than the IPCs. In the SVZ, Dlx2-(GABAergic phenotype) and Tbr2-(glutamatergic phenotype) expressing neuroblasts have been found (Lledo et al. 2008; Hodge et al. 2012). In the SGZ, glutamatergic Tbr2-expressing neuroblasts have been predominantly detected (Hodge et al. 2012), though dorsal pallial progenitors also generate a specific subset of GABAergic interneurons (Kohwi et al. 2007; Cai et al. 2013). The expression of neuronal lineage markers as the cytoskeletal proteins Doublecortin (DCX) and the polysialylated-neural cell adhesion molecule (PSA-NCAM) are usually used as markers of migratory neuroblasts. The absence of proliferation markers in these cells allows to differentiate postmitotic neuroblasts from progenitor cells (Hodge et al. 2012).

Adult neurogenesis has also been studied in the telencephalon of other non-mammalian species such as birds (Goldman and Nottebohm, 1983; Nordeen and Nordeen, 1988a, b; Álvarez-Buylla et al. 1990, 1992; 1998, Walton et al. 2012; Mazengenya et al. 2018), reptiles (Pérez-Cañellas and García-Verdugo, 1996; Font et al. 2001), amphibians (Simmons et al. 2008; Kirkham et al. 2014; Joven and Simon, 2018) and fish (Adolf et al. 2006; Grandel et al. 2006; Zupanc, 2006; März et al. 2010; Quintana-Urzainqui et al. 2015; reviewed by Ganz and Brand, 2016). Similarities between mammals and these groups have been found regarding the main types of cells found in the telencephalic neurogenic niche and the molecular markers they express. However, their organization within the niche differs among vertebrates. In the SVZ of mammals, B, C and A cells are located subventricularly lining the ventricular zone (VZ), an epithelial monolayer of non-proliferative cuboid ependymal cells that separates the SVZ from the ventricular cavity (García-Verdugo et al. 2002). In reptiles and birds, radial glia-like progenitors (B cells) and migrating precursors (A cells) have been found within the VZ (usually a pseudostratified epithelium up to four nuclei deep) intermingled with radial ependymal cells (García-Verdugo et al. 2002), but transit amplifying or intermediate progenitors (C cells) have not been described to date. In zebrafish, dividing radial glia, intermediate progenitor cells and neuroblasts are found in the VZ of the telencephalon (März et al. 2010; Than-Trong and Bally-Cuif, 2015), which mostly lacks ependymal-like cells.

While in mammals the distinction between progenitor radial glia and differentiated glia is clear, in fish, but also in amphibians and reptiles, the term radial glia has been used to refer both to radial glial progenitors and to radial ependymoglia, which is the main glial cell type present in the mature brain of fish and amphibians (reviewed by Cuoghi and Mola, 2009; Allen and Lyons, 2018). Although progenitor radial glial cells and radial ependymoglia are cells with different degree of differentiation, in anamniotes both cell types express similar astroglial markers (Kirkham et al. 2014; Than-Trong and Bally-Cuif, 2015).

Adult neurogenesis in different species has been suggested to be related to regenerative capacity, learning, spatial, contextual and emotional memories (Augusto-Oliveira et al. 2019 and references therein). Comparative studies have evidenced that the neurogenic capacity in the adult becomes more restricted to anterior regions of the brain throughout the course of vertebrate evolution. Fish are the group of vertebrates with the highest neurogenic potential, which has been linked to a continuous growth of the brain and with a high regenerative capacity (Alunni and Bally-Cuif, 2016; Ganz and Brand, 2016). However, most studies in fish have been carried out in modern teleost fish as zebrafish, and almost none have been carried out in cartilaginous fish (Quintana-Urzainqui et al. 2015).

The telencephalon of cartilaginous fish is a large non-layered structure that represents the 50% of the total cerebral mass (Yopak et al. 2015). Contrary to the everted telencephalon of teleost (Nieuwenhuys, 2009), the telencephalon of cartilaginous fish develops by evagination as in all other jawed vertebrates, which eases comparative studies. Besides, their phylogenetic position as a sister group of gnathostomes with a bony skeleton that gave rise to land vertebrates makes them essential in assessing the ancestral condition of particular traits in the brain of jawed vertebrates (Rodríguez-Moldes et al. 2017). Developmental studies in cartilaginous fish have evidenced high similarities to mammals concerning proliferating patterns and migratory routes in the developing telencephalon (Carrera et al. 2008; Quintana-Urzainqui et al. 2015; Docampo-Seara et al. 2018). Concerning adult neurogenesis, recent studies have evidenced the existence of abundant cells expressing proliferating markers in the ventricles leading to the olfactory bulbs and in restricted areas of the adult telencephalon. These regions also expressed GFAP and markers of migrating neuroblasts as DCX, which suggest the existence of neurogenic niches similar to those described in mammals (Quintana-Urzainqui et al. 2015). However, a deep molecular characterization of different types of progenitor cells and new-born neurons and their cell organization in the adult telencephalic niche of cartilaginous fishes is lacking.

With the aim of extending the knowledge on the evolution of adult neurogenesis, we have performed a detailed analysis of the proliferating niches in the telencephalon of juvenile and adult specimens of the lesser spotted dogfish *Scyliorhinus canicula* or catshark, by using antibodies against the proliferating cell nuclear antigen (PCNA). Then we have characterized different types of progenitor cells located in the neurogenic niches of the telencephalon. We have investigated the expression pattern of the stem cell factor Sox2 and the radial glial markers GFAP, BLBP and GS, typically used for detecting progenitor radial glial cells in vertebrates. We have also examined the expression of the neuronal lineage markers *ScDlx2*, *ScTbr2* and the expression of DCX in order to determinate the possible existence of IPCs and neuroblasts and how are they organized within the neurogenic niche.

## Materials and Methods

### Experimental animals

We analyzed 15 juveniles of *S. canicula* from 10 to 25 cm long (early and late juveniles) and 3 adult specimens (50 cm long). Individuals were kindly provided by the aquarium of O Grove (Galicia, Spain). Catsharks were raised in seawater tanks under standard conditions of temperature (15-16 °C), pH (7.5-8.5) and salinity (35 g/L); and suitable measures were taken to minimize animal pain and discomfort. All procedures conformed to the guidelines established by the European Communities Council Directive of 22 September 2010 (2010/63/UE) and by Spanish Royal Decree 1386/2018 for animal experimentation and were approved by the Ethics Committee of the University of Santiago de Compostela.

### Tissue processing

Juveniles were deeply anesthetized with 0.5% tricaine methane sulfonate (MS-222; Sigma, St. Louis, MO) in seawater and then perfused intracardially with elasmobranch Ringer’s solution (see Ferreiro-Galve et al. 2012) followed by 4 % PFA in Elasmobranch Phosphate Buffer (EPB). Brains were removed and postfixed in the same fixative for 24-48 h at 4 °C. Subsequently, they were rinsed in PB saline (PBS), cryoprotected with 30 % sucrose in PB, embedded in OCT compound (Tissue Tek, Torrance, CA), and frozen with liquid nitrogen-cooled isopentane. Parallel series of sections (18-20 μm thick) were obtained in transverse planes on a cryostat and mounted on Superfrost Plus (Menzel-Glasser, Madison, WI, USA) slides.

### In situ hybridization

We applied in situ hybridization (ISH) for *S. canicula Sox2, Tbr2/Eomes* and *Dlx2 (ScSox2, ScTbr2, and ScDlx2)* probes. These genes were selected from a collection of *S. canicula* embryonic cDNA library (mixed stages S9 to 22) and submitted to high throughput EST sequencing (coordinated by Dr. Sylvie Mazan). Sense and antisense digoxigenin-UTP-labeled *ScSox2, ScTbr2* and *ScDlx2* were synthesized directly by transcription *in vitro*. ISH was performed on cryostat sections of juveniles following standard protocols (Coolen et al. 2007). Briefly, sections were permeabilized with proteinase K, hybridized with sense or antisense probes overnight at 65° C and incubated with the alkaline phosphatase-coupled anti-digoxigenin antibody (1:2000, Roche Applied Science, Manheim, Germany) overnight at 4° C. The color reaction was performed in the presence of BM-Purple (Roche). Finally, sections were dehydrated and coverslipped. Control sense probes did not produce any detectable signal.

### Immunohistochemistry

Sections were pre-treated with 0.01 M citrate buffer pH 6.0 for 30 min at 90 °C for heat-induced epitope retrieval and allowed to cool for 20 min at room temperature (RT). Sections were rinsed in 0.05 M Tris-buffered saline (TBS) pH 7.4 for 5 min and treated with 10% H_2_O_2_ in TBS for 30 min at RT to block endogenous peroxidase activity. Sections were rinsed in 0.05 M TBS pH 7.4 for 5 min and incubated approximately for 15 h at RT with primary antibodies (see Table 1). Sections were rinsed three times in 0.05 M TBS pH 7.4 for 10 min each, and incubated in the appropriate secondary antibody (see Table 1) for 1 hour at RT. All dilutions were made with TBS containing 15% normal goat serum (Millipore, Billerica, MA), 0.2% Triton X-100 (Sigma) and 2% bovine serum albumin (BSA, Sigma). All incubations were carried out in a humid chamber. Then, sections were rinsed three times in 0.05 M TBS pH 7.4 for 10 min each. The immunoreaction was developed with 0.25 mg/ml diaminobenzidine (DAB) tetrahydrochloride (Sigma) in TBS pH 7.4 and 0.00075 % H_2_O_2_, or with SIGMAFAST™ 3.3-DAB tablets as indicated by the manufacturers. For immunohistochemistry against PCNA in adult individuals and PH3 in juveniles, 2.5 mg/mL of nickel ammonium sulphate was added. Finally, the sections were dehydrated, and coverslipped. Information about the primary and secondary antibodies is included in Table 1.

**Table 1.** Primary and secondary antibodies used in this study.

| **<u>PRIMARY ANTIBODY</u>** | **<u>SOURCE</u>** | **<u>WORKING DILUTION</u>** |
| --- | --- | --- |
| BLBP | Policlonal rabbit anti-BLBP<br>Millipore (Cat n°. ABN14) | 1:300 |
| BrdU | Monoclonal mouse anti-BrdU<br>Millipore (Cat n°. MAB4072) | 1:200 |
| DCX | Policlonal rabbit anti-DCX<br>Cell Signalling (Cat n°. 4604) | 1:300 |
| GAD | Policlonal rabbit anti-GAD<br>Millipore (Cat n°. AB1511) | 1:300 |
| GFAP | Policlonal rabbit anti-GFAP<br>Dako (Cat n°. Z0334) | 1:500 |
| GS | Monoclonal mouse anti-GS<br>Millipore (Cat n°. MAB302) | 1:500 |
| PCNA | Monoclonal mouse anti-PCNA<br>Sigma (Cat. n° P8825) | 1:500 |
| **<u>SECONDARY ANTIBODY</u>** | **<u>SOURCE</u>** | **<u>WORKING DILUTION</u>** |
| Goat anti-mouse HRP coupled | Dako, Glostrup, Denmark | 1:200 |
| 488-cojugated donkey anti-mouse | Alexa Fluor<br>Molecular Probes, Eugene, OR | 1:200 |
| 546-conjugated donkey anti-rabbit | Alexa Fluor<br>Molecular Probes, Eugene, OR | 1:200 |
| FITC-conjugated goat anti-mouse | ThermoFisher<br>Cat n°. F2761 | 1:100 |
| Cy3 conjugated goat anti-rabbit | ThermoFisher<br>Cat n°. A10520 | 1:200 |

### Double in situ hybridization-immunohistochemistry

We applied double ISH-immunohistochemistry for *ScSox2, ScTbr2 and ScDlx2* probes and PCNA antibody. In this procedure, ISH has been performed first, following the procedure described above. Color reaction was stopped by rinsing twice in phosphate buffer solution (PBS) for 10 min each and then in PFA 4% for 45 min. Then, immunohistochemistry was performed as described above.

### Double immunofluorescence

For heat-induced epitope retrieval, sections were pre-treated with 0.01 M citrate buffer (pH 6.0) for 30 min at 90 °C and allowed to cool for 20 min at RT. Sections were rinsed in 0.05 M TBS pH 7.4 for 5 min and incubated approximately for 15 h at RT with primary antibodies (see Table 1). Sections were rinsed three times in 0.05 M TBS pH 7.4 for 10 min each, and incubated in the appropriate combination of fluorescent dye-labelled secondary antibodies (see Table 1) for 1 hour at RT. All dilutions were made with TBS containing 15 % normal donkey serum (Millipore, Billerica, MA) 0.2 % Triton X-100 (Sigma) and 2 % BSA (Sigma). All incubations were carried out in a humid chamber. Sections were rinsed three times in 0.05 M TBS pH 7.4 for 10 min each and in distilled water for 30 min too. Sections were then allowed to dry for 30 min at 37 °C, and mounted in MOWIOL 4-88 Reagent (Calbiochem, MerkKGaA, Darmstadt, Germany). Information about the primary and secondary antibodies is included in Table 1. Eventually, nuclei were counterstained with blue-fluorescent DAPI nucleic acid stain (Vectashield mounting medium for fluorescence with DAPI; Vector, Burlingame, California).

### BrdU experiments

BrdU pulse-chase labelling experiments were performed by incubating 1 catshark juvenile (13 cm long) with 5 mg/ml of BrdU in oxygenated artificial sea water for 24 h. Then it was anesthetized and sacrificed by intracardiac perfusion with Ringer solution and PFA 4% and postfixed by immersion in PFA 4% for 48 h. For detection of BrdU, sections were incubated in 2N HCl for 30 min at 50 °C to denature DNA strands. HCl reaction was stopped by addition of 0.1 M sodium tetraborate and sections were then rinsed in TBS for 10 min before antibody incubation. Sections were incubated with the proper anti-BrdU antibody (see Table 1) at RT overnight and processed for immunofluorescence as described above.

### Control and specificity of antibodies

The PCNA antibody has been previously used to label progenitor cells in the brain and retina of catshark (i.e. Quintana-Urzainqui et al. 2015; Sánchez-Farías and Candal, 2015). The anti-phosphohistone 3 (PH3) antibody has been previously used as a marker of mitotic cells in the telencephalon of catshark (Quintana-Urzainqui et al. 2015; Docampo-Seara et al. 2019). The specificity of the antibodies against the glial markers GFAP, BLBP and GS has been tested by western blot (Docampo-Seara et al. 2019). The anti-GAD antibody used in this study shows the same expression pattern than the anti-GAD65/67 antibody previously used in the identification of subpallial-derived GABAergic cells in the developing pallium of the catshark (Carrera et al. 2008; Quintana-Urzainqui et al. 2015). The specificity of the anti-DCX antibody has been also tested by western blot by Pose-Méndez et al. (2014).

### Imaging

Fluorescent sections were photographed with the Leica TCS-SP2 scanning microscope with a combination of blue and green excitation lasers. Confocal images were acquired separately for each laser channel with steps of 1 μm along the z-axis, and collapsed images were obtained with the LITE software (Leica). On the other hand, light field images were obtained with an Olympus BX51 microscope equipped with an Olympus DP71 color digital camera. Both fluorescent and light field photographs were adjusted for contrast, brightness and intensity using Corel Draw X7. Plates also were prepared using the same software.

### Cell counting

Cell counting was performed using 5 juveniles (10-13 cm long) and 3 adults (50 cm long). The area selected was the VZ of the ventral pallium since it is the area that hosts higher numbers of proliferating cells per surface, both in juveniles and adults (see Fig. 1). In adults, cells showing either weak or intense immunoreactivity to PCNA were counted in a square box of 50×50 µm in two selected fields containing high density (HD) and low density (LD) of PCNA-immunoreactive (-ir) cells, respectively. In juveniles, cells were counted in two random 50×50 µm fields, since PCNA-ir cells are homogenously distributed. Cells were counted manually, and average and standard deviations were calculated using Microsoft Excel 2016.

**Figure 1.**
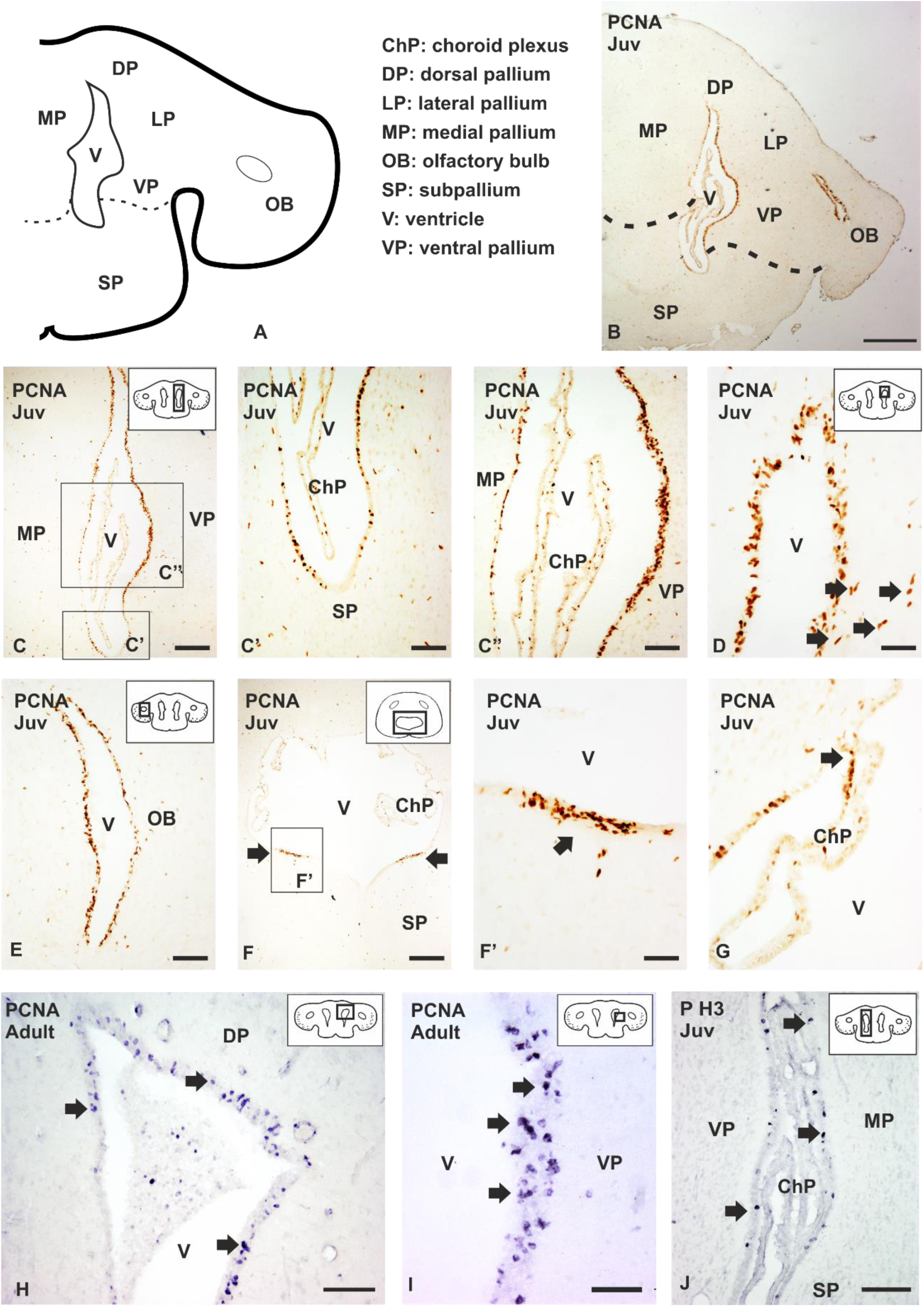
Anatomical scheme of a transverse section of the telencephalon of *Scyliorhinus canicula* **(A),** and photomicrographs from transverse sections at different magnifications showing the expression pattern of PCNA in juveniles (Juv) (**B-G),** adults **(H-I),** and the expression pattern of PH3 in juveniles **(J)** of *S. canicula*. **(B)** Panoramic view of the telencephalon of a juvenile showing PCNA immunoreactivity in the VZ of the lateral telencephalic ventricle and the ventricle of the olfactory bulb. **(C-C’’)** Photomicrographs at high magnifications of the VZ of the telencephalon showing different density of PCNA immunoreactive cells among the subpallium **(C-C’)** and the different subdivisions of the pallium **(C, C’’).** Note that the pallium shows a higher density of PCNA-ir cells than the subpallium. **(D)** Photomicrograph at high magnification of the dorsal pallium showing PCNA immunoreactive (PCNA-ir) located lining the ventricle and some immunoreactive cells located subventricularly (arrows). **(E)** Photomicrograph at high magnification of the olfactory bulb showing numerous PCNA-ir cells in the ventricular zone. **(F-G)** Photomicrographs at different magnifications of the caudal telencephalon showing numerous PCNA-ir cells restricted to the lateral regions of the subpallial VZ (arrows) **(F-F’)** and cells that contain PCNA in the choroid plexus (arrow in **G**). **(H-I)** Photomicrographs at different magnifications of an adult brain showing PCNA-ir cells in the dorsal **(H)** and ventral **(I)** pallium (arrows). **(J)** Photomicrograph of the adult telencephalon showing scattered PH3 immunoreactive cells in the ventricular zone of the lateral telencephalic ventricle (arrows). Dotted lines in the anatomical scheme represent the pallial-subpallial boundary. Scale bars: 500 µm (B) 200 µm (C, F, H), 100 µm (C’, C’’, E, I), 50 µm (D, F’, G, J).

## Results

The telencephalon of *S. canicula* has been classically subdivided in three parts: the olfactory bulbs, the telencephalic hemispheres (pair ventricles, divided in pallium and subpallium) and the impar or caudal telencephalon (enclosing an impar ventricle, from the anterior commissure to the caudal part of the optic chiasm; Smeets et al. 1983). In this classic view, the telencephalic hemispheres are located topographically rostral to the impar telencephalon that, in turn, is located rostral to the hypothalamic subdivisions (See Supp. Fig. 1A-C). However, modern neuroanatomy evidences that this view does not reflect the topologically invariant organization of the telencephalon (Nieuwenhuys and Puelles, 2016). According to the prosomeric model, the prosencephalon is divided in two transverse segments termed prosomeres (Supp. Fig. 1D, E). The caudal one (hp1) includes the peduncular hypothalamus and the evaginated bilateral telencephalic vesicles (divided in pallium and evaginated subpallium). The rostral one (hp2) includes the terminal hypothalamus and the preoptic area, a non-evaginated subpallial compartment located between the anterior commissure and the hypothalamus (Nieuwenhuys and Puelles, 2016; for more information about the prosomeric model in the catshark see Santos-Durán et al. 2015 and Rodríguez-Moldes et al. 2017). Despite developmental and genoarchitecture studies in *S. canicula* support the prosomeric model, descriptions in what follows are made according to rostro-caudal topographical axes that is the prevailing way to describe neuroanatomical subdivisions in the adult catshark.

### Proliferation pattern

In the present study we have investigated the expression pattern of the proliferating cell nuclear antigen (PCNA) in order to determine the presence of cell proliferation in the telencephalon of juveniles and adults of *S. canicula.* PCNA is present in the interphase of the cell cycle in proliferating cells, and its expression is stronger during the S phase (Zerjatke et al. 2017). In juveniles PCNA-ir cells are mainly located in the VZ of the olfactory bulb, lateral ventricles of the telencephalic hemispheres and in the VZ of the impar telencephalon. Rostrally, in the telencephalic hemispheres, PCNA-ir cells are distributed homogeneously through the VZ of the pallial and subpallial subdivisions (not shown). In intermediate levels of the telencephalon (Fig. 1A, B), numerous PCNA-ir cells are also appreciated, but remarkable differences among different telencephalic regions are observed. First, PCNA positive cells are considerably more abundant in the VZ of the pallium than in the subpallium (Fig. 1B-C’’). In addition, differences in the density of PCNA-ir cells are observed along the pallial VZ: the dorsal, lateral and ventral regions of the pallial subdivision exhibit considerably more PCNA-ir cells than the medial pallial subdivision (Fig. 1B, C, C’’). Also, fusiform PCNA-ir cells can be detected close to the dorsal pallial VZ (arrows in Fig. 1D). The VZ of the olfactory bulbs have also numerous PCNA-ir cells (Fig. 1E). In the impar telencephalon (Fig. 1F), numerous PCNA-ir cells are observed in the lateral portions of ventricular surface (arrows in Fig. 1F, F’), but the medial region is devoid of PCNA immunoreactivity. Disperse adventricular PCNA-immunoreactive (-ir) cells are detected both in the pallium and subpallium, as well as in the olfactory bulb (Fig. 1B-E), and their amount decreases in the impar telencephalon (Fig. 1F, F’). In addition, immunoreactive cells can be appreciated in the choroid plexus, specially concentrated in the region that contacts the VZ (arrow in Fig. 1G). Similar patterns of proliferation can be observed in adult specimens (50 cm long; Fig. 1H-I).

Since PCNA expression can be detected not only during the interphase in proliferating cells but also for a few hours after induced cell quiescence (Zerjatke et al. 2017), we have additionally investigated the presence of dividing cells by analyzing the expression of PH3, a mitosis marker. As expected, PH3-ir cells are found along the VZ of the telencephalic hemispheres and they are more numerous in the pallium than in the subpallium (arrows in Fig. 1J).

In order to evaluate possible variations in proliferation rates as maturation progresses, PCNA-ir cells were counted within the VZ of the ventral pallium both in juveniles and adults. We selected the VZ of the ventral pallium due to it exhibits high numbers of PCNA-ir cells. While the density of PCNA positive cells per surface decreases in adult specimens, there are not significative differences in the absolute number of PCNA-ir cells in the VZ between juveniles and adults (see Supp. Fig. 2) and therefore we have used juvenile specimens in what follows.

**Figure 2.**
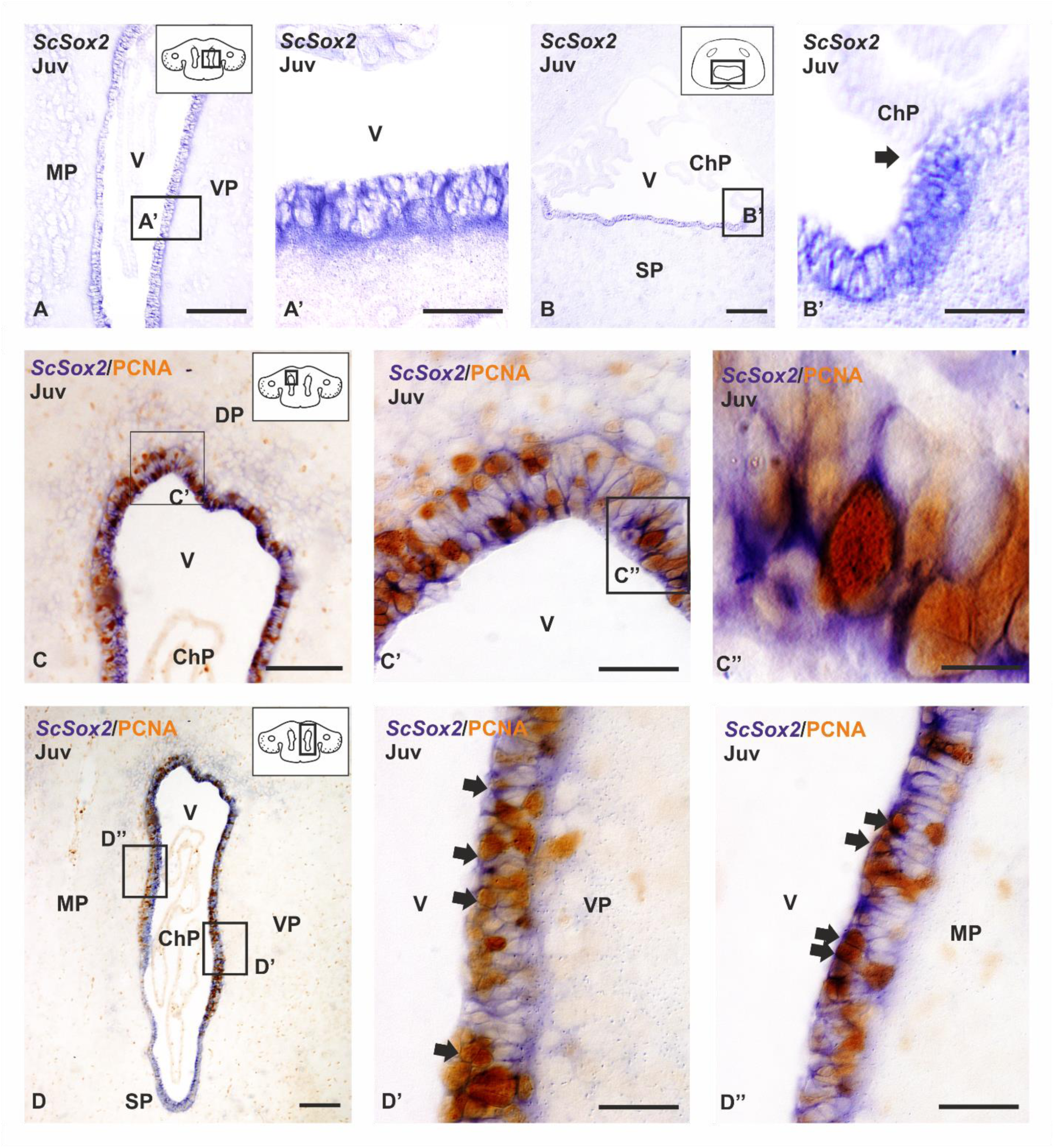
Photomicrographs at different magnifications from transverse sections showing the expression pattern of *ScSox2* **(A-B’)** and *ScSox2-*PCNA immunoreactivity **(C-D’’)** in the telencephalon of catshark juveniles (Juv). **(A-B’)** Photomicrographs at different magnifications showing the high *ScSox2* expression in the VZ of the lateral telencephalic ventricles (**A, A’**) and caudal telencephalon (**B, B’**). Note that in the caudal telencephalon *ScSox2* is not expressed in the choroid plexus (**arrow in B’**). **(C-D’’)** Photomicrographs at different magnifications showing double ISH-IHC between *ScSox2-*PCNA in the dorsal **(C-C’’),** ventral **(D-D’)** and medial pallium **(D, D’’)** (arrows). Note differences between the medial and the ventral pallium regarding the thickness of the VZ and coexpression of *ScSox2* and PCNA **(D-D’’).** Scale bars: 200 µm (B, D), 100 µm (A, C), 50 µm (A’, B’, C’, D’, D’’), 10 µm (C’’). Abbreviations: ChP, choroid plexus; DP, dorsal pallium; MP, medial pallium; SP: subpallium; V, ventricle; VP: ventral pallium.

### Sox2 expression pattern

In the adult mammalian brain, two subpopulations of progenitor cells (radial glia and early IPCs) retain the expression of the stem cell marker Sox2 (Bonaguidi et al. 2016). In addition, previous studies in adult zebrafish have reported that, in the VZ of the telencephalic hemispheres, most ventricular cells express Sox2 (März et al. 2010). In order to investigate whether Sox2 is expressed in the whole ventricular layer or it defines subpopulations of progenitor cells as in mammals, we studied the expression pattern of *ScSox2* by *in situ* hybridization in the telencephalon of *S. canicula*.

In juveniles, most cells in the VZ of the olfactory bulbs (not shown) and the telencephalic hemispheres (Figs. 2A, A’) are positive for *ScSox2*. Also, *ScSox2* expressing cells are present at the level of the impar ventricular surface (Fig. 2B). *ScSox2* expressing cells were absent from the choroid plexus (arrow in Fig. 2B, B’).

Since Sox2 is a marker of progenitor cells (see above) and the presence of PCNA is commonly used to distinguish proliferating from quiescent or post-mitotic cells in fixed samples (reviewed in Zerjatke et al. 2017), we then performed double ISH-immunohistochemistry for *ScSox2* and PCNA. Numerous double-labeled cells for *ScSox2* and PCNA can be observed in the olfactory ventricular surface (not shown) and in the VZ of all pallial areas studied, including the dorsal and lateral pallium (Figs. 2C-C’’), the ventral pallium (Figs. 2D, D’, arrows) and the medial pallium (Figs. 2D, D’’, arrows). A few double labeled cells are also observed in the subpallium (not shown). However, numerous *ScSox2*-expressing cells were PCNA-negative in the pallium.

### Double immunofluorescence against glial markers

Studies in the adult zebrafish have established that, in addition to Sox2, most progenitor cells in the VZ express glial markers as GFAP, BLBP and GS, though some of these glial markers are also expressed by non-progenitor glial cells including ependymal cells (reviewed by Than-Trong and Bally-Cuif, 2015). Most cells in the telencephalic VZ of the catshark express *ScSox2* (see above) and numerous GFAP-, BLBP-, and GS-expressing radial ependymoglia have been previously described lining the telencephalic ventricles in early juveniles of catshark (Docampo-Seara et al. 2019). In order to see if in the catshark, in contrast to zebrafish, the combined expression of these markers allows to differentiate different types of progenitor cells from non-progenitor cells, we have performed double immunofluorescence against the radial glial markers GS/GFAP and GS/BLBP. Then, we have combined them with PCNA in order to know if they correspond to quiescent or dividing cells. We have mainly focused our analysis in the medial pallium and in the ventral pallium as representative areas containing low and high densities of proliferating cells, respectively.

Double immunofluorescence against GS and GFAP shows that the vast majority of cells are double-labelled (Fig. 3A-C’’). Positive cells expressing only GS or GFAP are scarce (stars in Figs. 3C-C’’) and are located close to each other. On the other hand, double immunofluorescence against GS and BLBP shows that both molecular markers colocalize in the same cells (Figs. 3D-F’’), and their expression occurs in the same cell domains (Figs. 3F-F’’).

**Figure 3.**
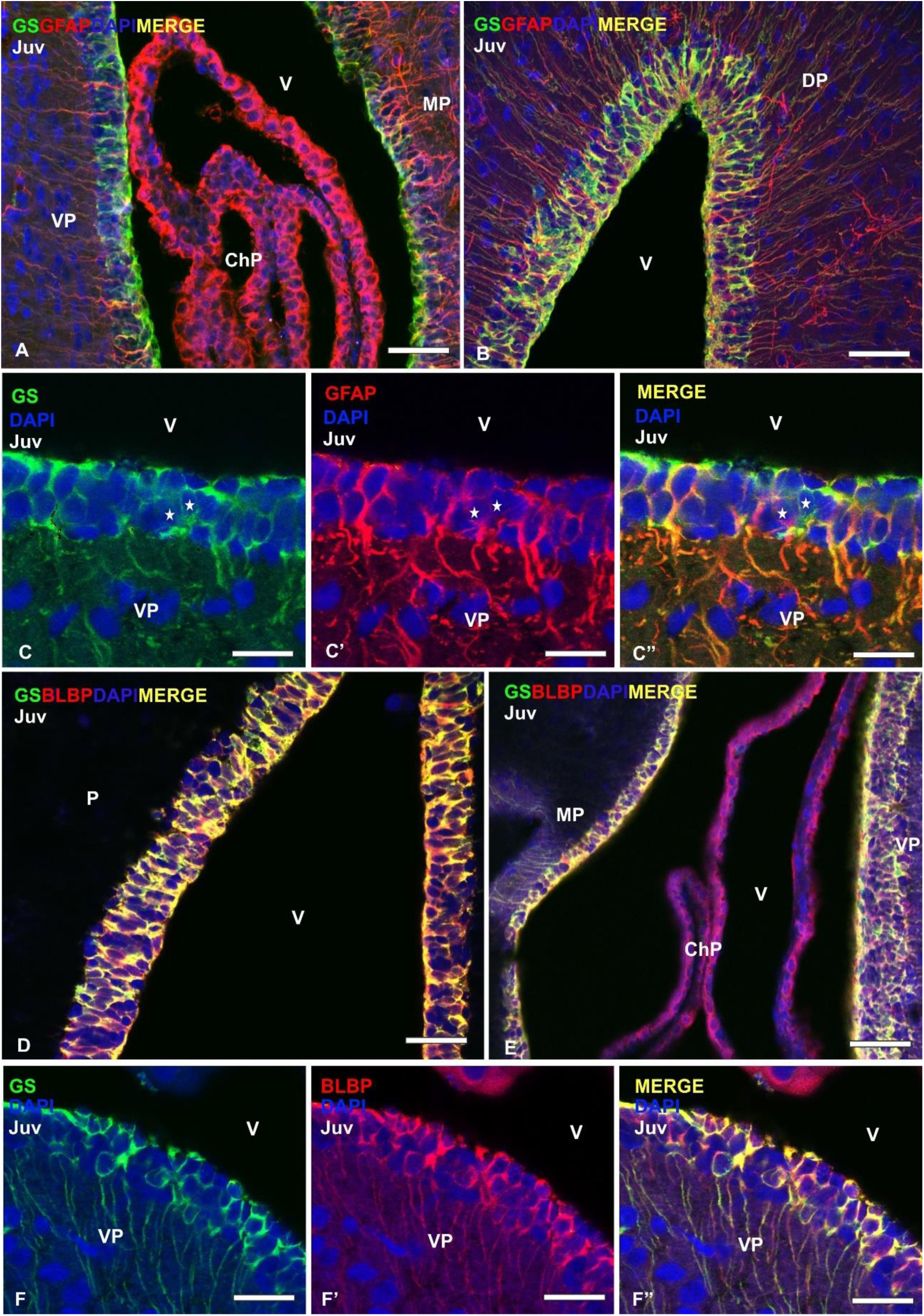
Photomicrographs at different magnifications from transverse sections of the telencephalon of juveniles (Juv) showing double immunofluorescence between GS/GFAP **(A-C’)** and GS/BLBP **(D-F’’)** counterstained with DAPI. **(A-B).** Photomicrographs at low (A) and high magnification (B) of the dorsal telencephalon showing numerous double labelled cells in the pallial ventricular zone. Note the high amount of immunoreactive processes that are arranged radially **(C-C’’).** Photomicrographs at higher magnification of the VZ of the ventral pallium showing the high degree of colocalization between GS and GFAP. Note also that a few cells are only positive for one of the glial markers (stars). Photomicrographs at low **(D-E)** and high magnification **(F-F’’)** of the ventral pallium VZ showing numerous cells where BLBP and GS are coexpressed. Scale bars: 100 µm (A, B, D, E), 25 µm C, C’, C’’, F, F’, F’’). Abbreviations: Chp, choroid plexus; DP, dorsal pallium; MP, medial pallium; P, pallium; VP: ventral pallium; v, ventricle.

Then, we have proceeded to combine glial markers with the proliferating marker PCNA. Double immunofluorescence against GFAP and PCNA revealed numerous double-labelled cells in the VZ of the telencephalon. In the rostral telencephalon, where no differences regarding PCNA immunoreactivity were observed along the VZ, some double-labeled cells have been observed (stars in Figs. 4A-A’’). However, in medial levels of the telencephalon, where the VZ of different telencephalic regions exhibits different levels of proliferation, the number of double-labelled cells increases considerably, especially in the ventral pallium, where the VZ seems to be wider than in other pallial regions of the telencephalic ventricle (Figs. 4B-B’’). Concerning BLBP, some double labeled cells can be appreciated in rostral levels, where the VZ show similar thickness. However, at intermediate levels, both in the medial and ventral pallium numerous double-labeled cells have been found (stars in Figs. 4C-C’’ and 4D-D’’ respectively).

**Figure 4.**
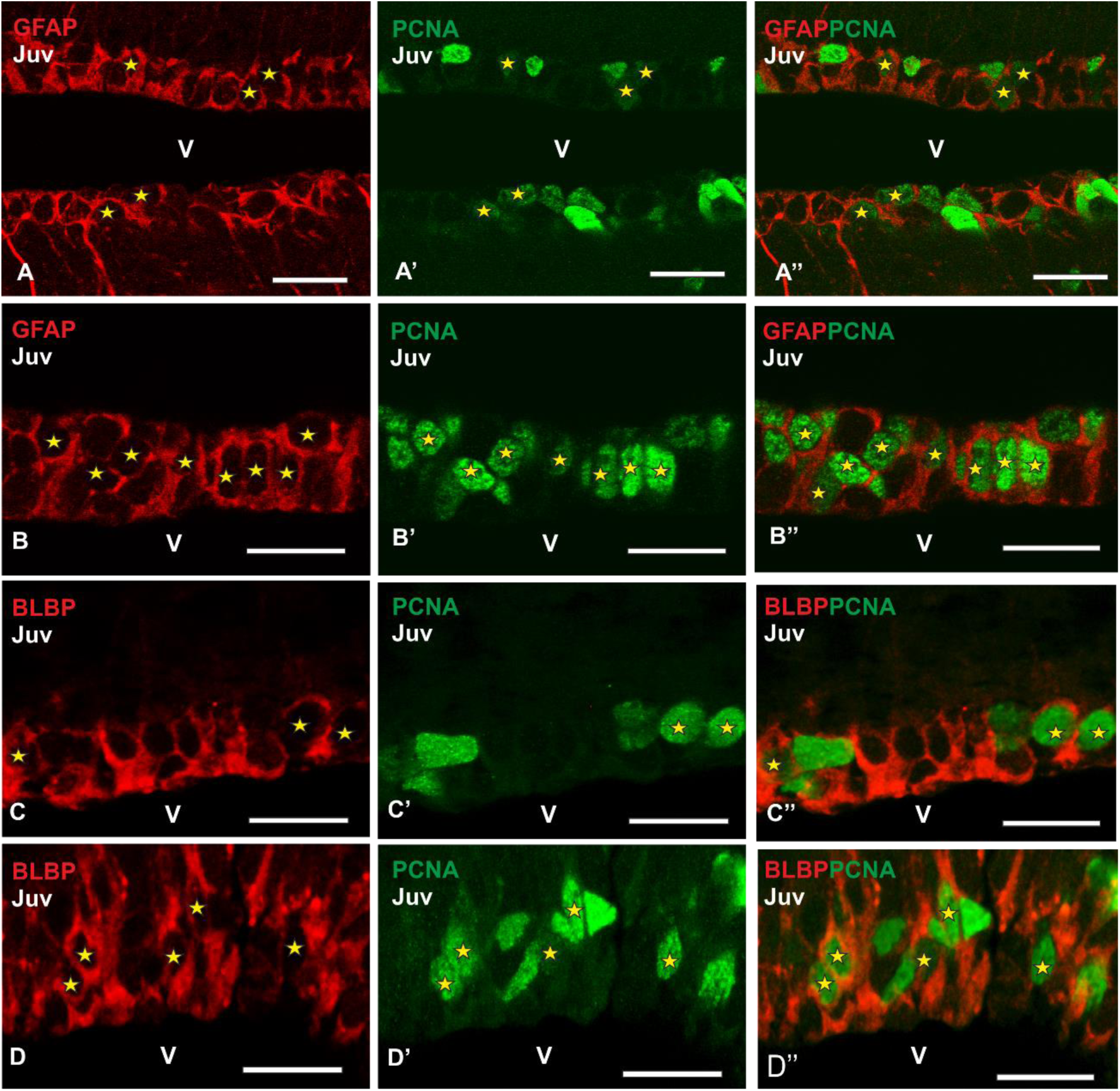
Photomicrographs at high magnification from telencephalic transverse sections of catshark juveniles (Juv) showing the immunoreactivity pattern of GFAP/PCNA **(A-B’’)** and BLBP/PCNA **(C-D’’)** in the lateral telencephalic ventricles. Details of the VZ of the rostral pallium **(A-A’’)** and ventral pallium **(B-B’’)** showing numerous double-immunolabeled cells for GFAP and PCNA (yellow stars). Photomicrographs at high magnification of the dorsomedial pallial VZ **(C-C’’)** and ventral pallium **(D-D’’)** where numerous cells coexpressing BLBP and PCNA are present (yellow stars). Scale bars: 25 µm. Abbreviations: v, ventricle.

### Expression of molecular markers of IPCs: Dlx2 *and* Tbr2

During development, two different genetic networks act early and spatially separate from each other in the mammalian and zebrafish embryo to generate excitatory (glutamate) or inhibitory (GABA) cells in the pallium and subpallium, respectively (Wullimann, 2009). Subpallial GABAergic precursors first migrate along spatially distinct tangential routes into the developing cortex, followed by the final radial migration into distinct cortical layers (Achim et al. 2014). During adult neurogenesis, in mammals, the neurogenic niches generate GABAergic and glutamatergic new neurons (Hevner et al. 2006; Lledo et al. 2008; Mind and Song, 2011; Hodge et al. 2012; Bond et al. 2015; Lim and Álvarez-Buylla, 2016). Progenitors in the SVZ (subpallium) produce multiple lineages of new neurons that include dopaminergic, GABAergic (i.e. Lledo et al. 2008), and a recently described subset of glutamatergic neurons, all of which migrate through the rostral migratory stream (RMS) to populate several areas of the olfactory bulb (Hodge et al. 2012). Progenitors in the SGZ (pallium) mainly produce glutamatergic projection neurons. Noteworthy, while the majority or pallial GABAergic interneurons in rodents are born in the subpallium and migrate tangentially to the pallium as described above, dorsal pallial progenitors also generate a specific subset of GABAergic interneurons that migrate to the olfactory bulb (Kohwi et al. 2007; Cai et al. 2013). These GABAergic and glutamatergic cell lineages are generated by intermediate progenitors (IPC) in the neurogenic niche, which predominantly express Dlx2 and Tbr2 respectively.

Here we have investigated whether PCNA-ir cells express markers of IPCs and what cell lineage they generate by studying the combined expression pattern of PCNA/*ScTbr2* and PCNA/*ScDlx2*. *ScTbr2* expressing cells can be observed in the olfactory bulb, dorsal, lateral and ventral pallial subdivisions but only in the ventral pallium they are located close to the VZ (Fig. 5A). *ScTbr2*-expressing cells were not observed in the medial pallium or in the subpallium. Double *ScTbr2*-expressing and PCNA-ir cells have not been observed in the telencephalic VZ (Fig. 5A’), though some double PCNA/ScTbr2-expressing cells can be observed out of the VZ (arrow in Fig. 5A’). *ScDlx2* expression in the telencephalon has been observed in the olfactory bulb, dorsal and medial pallium. In the subpallium, *ScDlx2* is expressed in the basal superficial area (a derivative of the lateral ganglionic eminence homologue in catshark; Fig. 5B). In the ventricular zone of the dorsal and medial pallium *ScDlx2* numerous PCNA-ir cells are located (see Fig. 5B’) and scattered proliferating cells are also present in subventricular positions (arrowheads in Fig. 5B’). Double-labeled *ScDlx2*-PCNAcells are clearly observed in the VZ (Figs. 5C, C’, D, arrows).

**Figure 5.**
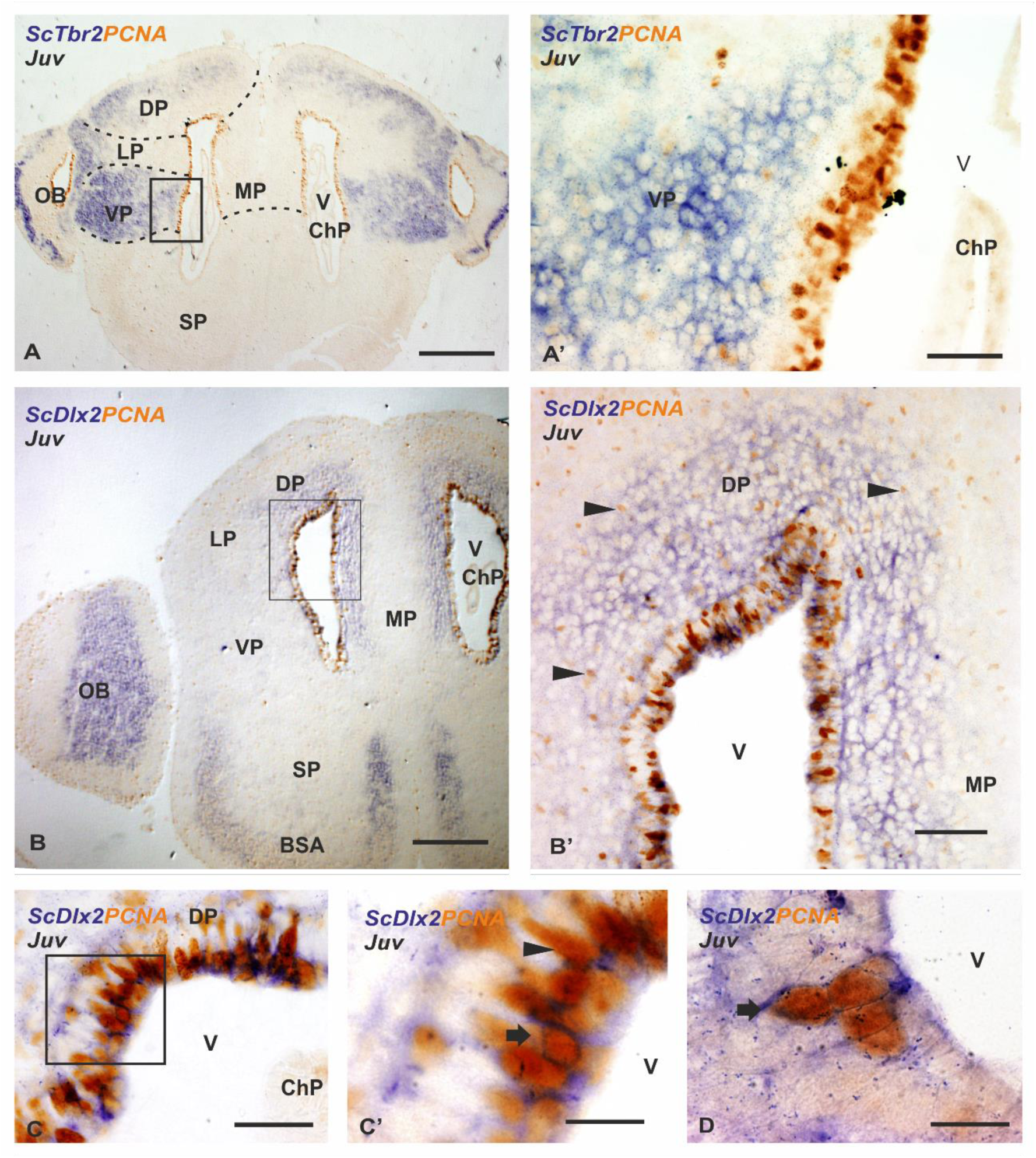
Photomicrographs at different magnifications from transverse sections showing the expression pattern of *ScTbr2-*PCNA **(A-A’)** and *ScDlx2-*PCNA **(B-D)** in the telencephalon of catshark juveniles (Juv). **(A-A’)** Photomicrographs at low and high magnification, respectively, of the medial telencephalon to shown that *ScTbr2* is not expressed in the VZ of the telencephalic hemispheres where numerous PCNA-ir cells are present. *ScTbr2* is highly expressed out of the VZ in the ventral pallium, where some ScTbr2 expressing cells are also PCNA-ir (arrow in A’), in the dorsal and lateral pallium, and in the olfactory bulb. **(B-D).** Panoramic view **(B)** and photomicrographs at high magnifications **(B’’-D)** showing the expression pattern of *ScDlx2-*PCNA in the telencephalon. *ScDlx2* is highly expressed in the olfactory bulb, in the VZ of the lateral, dorsal and medial pallium, and in the basal superficial **(B)**. Note that most of the cells located away from the ventricle only express PCNA (arrowheads in B’’). Numerous *ScDlx2* positive cells coexpress PCNA in the ventricular zone (arrows in C’ and D) and a few cells expressing only PCNA are present (arrowheads in C’). Scale bars: 500 µm (A, B), 100 µm (B’), 50 µm (A’, C), and 10 µm (C, D). Abbreviations: BSA, basal superficial area; ChP, chohoid plexus; DP, dorsal pallium; LP, lateral pallium; MP, medial pallium; OB, olfactory bulb; SP: subpallium; v, ventricle; VP: ventral pallium.

Since a subset of GABAergic (Dlx2) cells in rodents are born in the pallium (see above), we performed a BrdU birthdating assay to investigate if pallial progenitors in *S. canicula* also give rise to cells of the GABAergic lineage (Supp. Fig. 3). Double BrdU/GAD-ir cells are observed in the pallial VZ (Supp. Fig. 3A-A’’). However, GABAergic progenitors are not present in the subpallium in juvenile specimens (Supp. Fig. 3B-B’’).

### Expression of a molecular marker of neuroblasts: DCX

As part of the neurogenic niche, a population of migratory neuroblasts have been found in mammals and other species of vertebrates (i.e Álvarez-Buylla et al. 1998; März et al. 2010; Ming and Song, 2011; Kirham et al. 2014; Bond et al. 2015; Macedo-Lima et al. 2016). For that, we have also investigated the distribution pattern of the microtubule associated protein DCX, a marker of migratory neuroblasts (Gleeson et al. 1999). DCX-immunoreactive cells were observed in both the VZ of the olfactory bulb and telencephalic hemispheres (Figs. 6A-A’’). DCX-immunoreactive cells are also observed subventricularly. No immunoreactivity for DCX has been found in the caudal VZ of the telencephalon (Fig. 6B). In order to further characterize DCX positive cells, we have performed double immunofluorescence for DCX and PCNA. We have observed abundant double-labelled cells in the VZ of pallium (yellow stars in Figs. 6C-C’’). In addition, DCX/PCNA-ir cells are also present at subventricular positions of the pallium (yellow stars in Fig. 6D-D’’).

**Figure 6.**
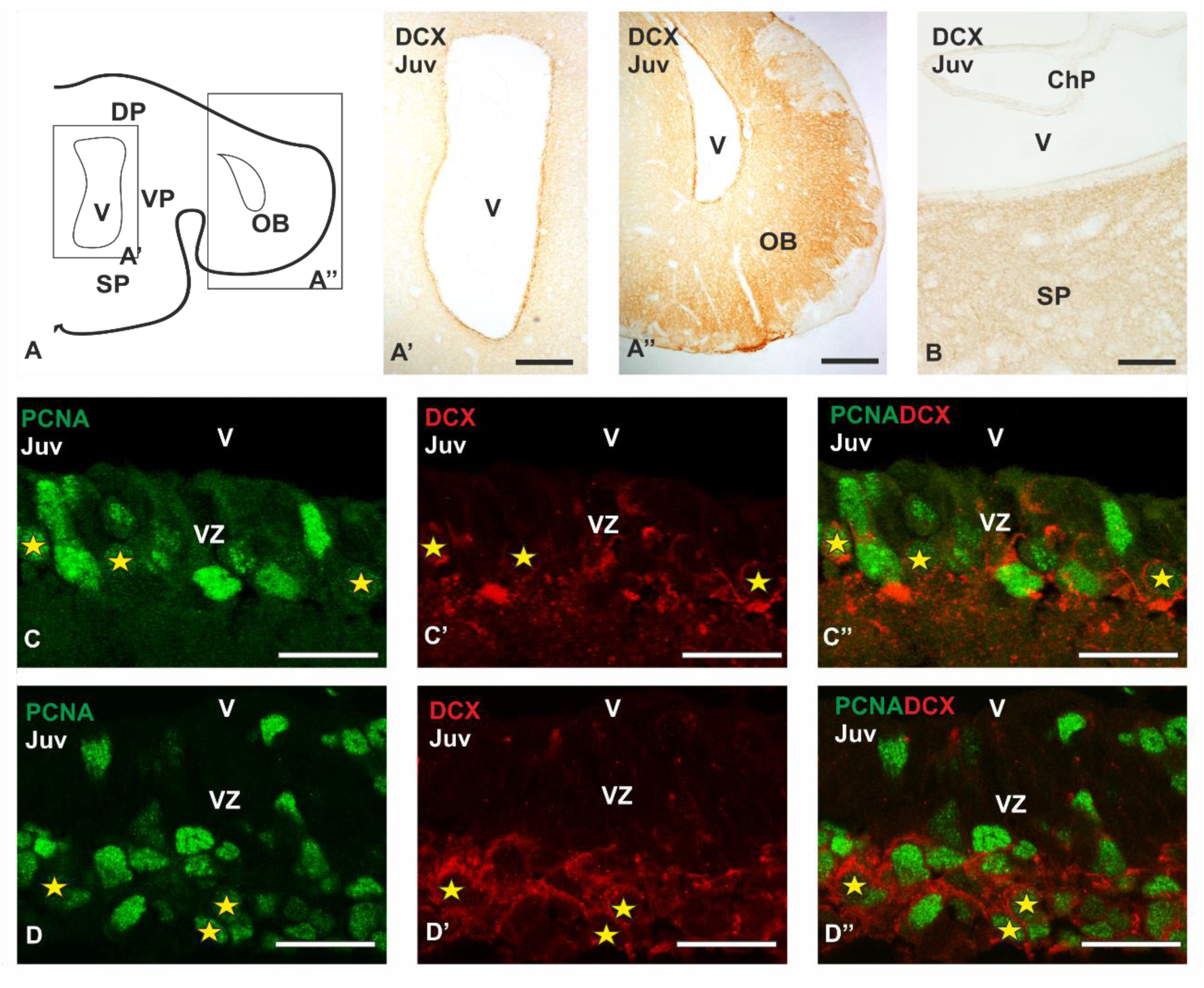
Anatomical scheme of a transverse section of the telencephalon of *Scyliorhinus canicula* **(A)** and photomicrographs at different magnifications from transverse sections of juveniles (Juv) processed by immunohistochemistry **(A-B)** and double immunofluorescence **(C-D’’)** showing the expression of DCX and coexpression of DCX-PCNA, respectively. **(A’-B)** Photomicrographs at low magnification of the lateral telencephalic ventricle **(A’)**, olfactory bulb **(A’’)** and caudal telencephalon **(B)** showing high DCX expression in these regions. Note that in the caudal telencephalon DCX is not expressed in the ventricular zone (B). **(C-D’’)** Photomicrographs at high magnification of the VZ **(C-C’’)** and territories close to the VZ **(D-D’’)** of the dorsal pallium showing numerous DCX-PCNA double-labelled cells **(yellow stars**). Scale bars: 200 µm (A’, A’’), 100 µm (B), 25 µm (C-D’’). Abbreviations: ChP, choroid plexus; DP: dorsal pallium; OB: olfactory bulb; SP: subpallium; VP: ventral pallium; V, ventricle; VZ: ventricular zone.

## Discussion

The study of adult neurogenesis has been a subject of interest during the past decades, not only in mammals, but also in many different species of birds (reviewed by Barnea and Pravosudov, 2011), reptiles (reviewed by González-Granero et al. 2011), amphibians (Simmons et al. 2008; Kirkham et al. 2014) and teleost fish (reviewed by Ganz and Brand, 2016). Previous reports from our group have shown that the telencephalon of sharks contains proliferative cells in the VZ of the adult telencephalon (Quintana-Urzainqui et al. 2015). With the aim of extending the knowledge about adult neurogenesis in the telencephalon of sharks we have analyzed the proliferating VZ of this species searching for different types of progenitor cells and studying their neuronal commitment. Then we have compared the molecular characteristics and organization of cells within the neurogenic niche of sharks with that of other vertebrates. A summary of the main cell types located in the neurogenic niches across vertebrates is provided in Figure 7.

**Figure 7.**
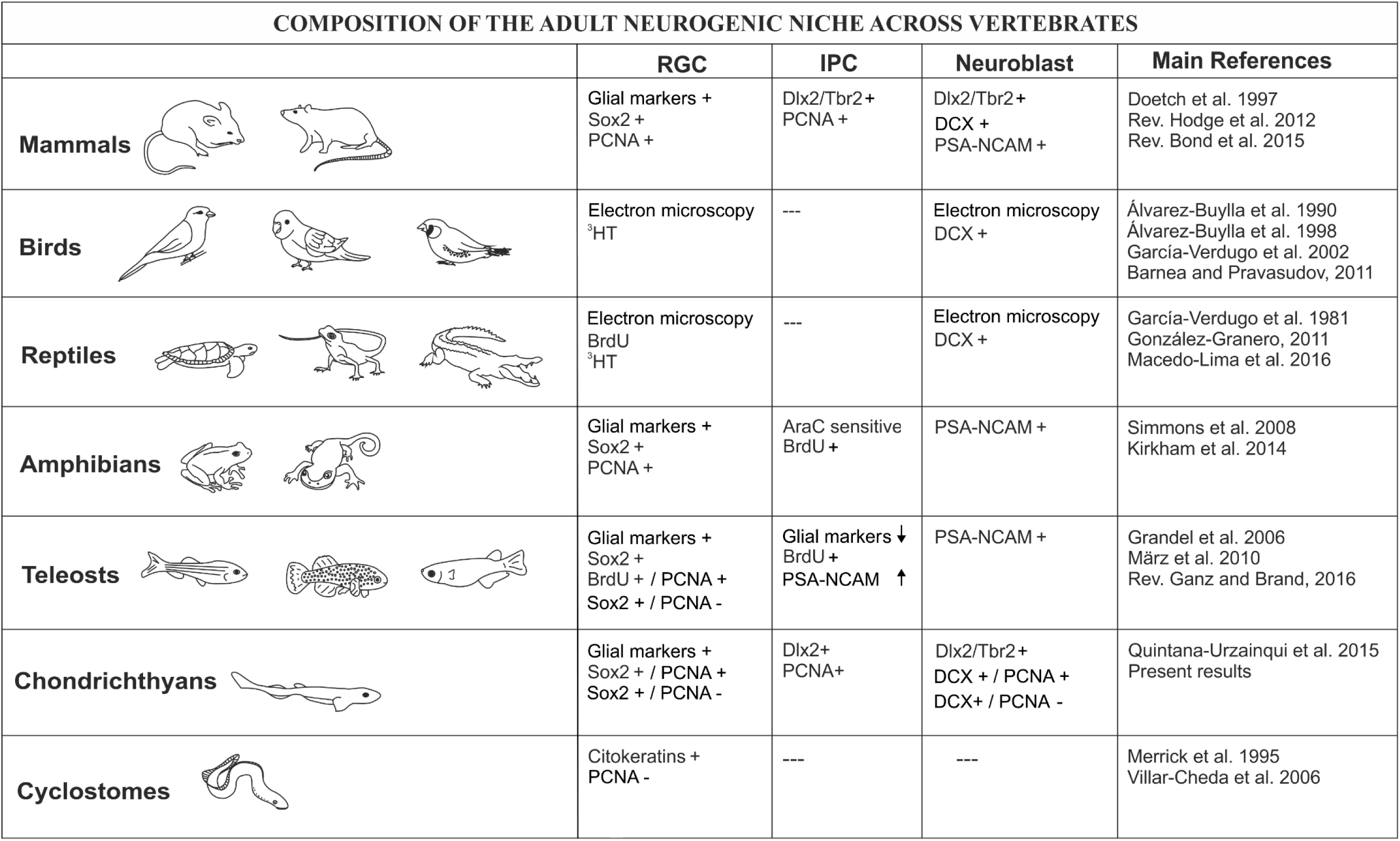
Summary of the main cell types involved in adult neurogenesis across vertebrate evolution.

### Telencephalic proliferation across evolution: the origin of adult neurogenesis

One of the most interesting facts of the neurogenic process is that the number of neurogenic niches and proliferating cells decreases considerably throughout evolution (Ganz and Brand, 2016). In mammals, neurogenesis is mainly restricted to some regions of the telencephalon such as the SVZ of the lateral ventricles and the SGZ of the dentate gyrus of the hippocampus (reviewed by Bond et al. 2015). In birds, adult neurogenesis has been detected in all major subdivisions of the telencephalon including the high vocal center (HVC; Goldman and Nottebohm, 1983) or the hippocampus (Barnea and Nottebohm, 1994) that display conspicuous regional differences in cell organization within the niche and in neurogenic capacity (reviewed in García-Verdugo et al. 2002). While the HVC is overlaid by a relatively thin VZ, in the other regions, dividing cells in the VZ tend to cluster in the called “hot spots”, a pseudostratified epithelium which contains diverse cell types. These “hot spots” coincide with areas of the VZ that are rich in radial glial cells and are located close to regions where young migrating neurons are firstly seen (Álvarez-Buylla et al. 1988a, b). In reptiles (mainly lizards) and amphibians (frog and salamander) the telencephalic VZ, exhibits proliferating cells (reptiles: reviewed by González-Granero et al. 2011; amphibians: Simmons et al. 2008; D’Amico et al. 2011; Kirkham et al. 2014). Despite proliferating cells have been found in all the pallial subdivisions and in the subpallium, these cells are not abundant. However, in amphibians, there are several “hot spots” (Álvarez-Buylla, et al. 1990; Kirkham et al. 2014). In adult teleost fish, the proliferation pattern has been extensively studied in the telencephalon of several species like zebrafish, medaka or killyfish, among others (Ekström et al. 2001; Candal et al. 2005; Zupanc et al. 2005; Grandel et al. 2006, Kuroyanagi et al. 2010; Tozzini et al. 2012; reviewed by Ganz and Brand, 2016). In this group of fish, the number of proliferative cells is drastically higher compared to tetrapod vertebrates and are usually observed as a continuous band of cells rather than being scattered in the VZ.

In the present study, we have investigated the proliferative cell pattern in the mature telencephalon of *S. canicula* by using antibodies against PCNA, which labels proliferating cells (reviewed in Zerjatke et al. 2017). As in teleost fishes, in *S. canicula* the paliall and subpallial VZ exhibit a rather continuous band of proliferating cells, though the subpallium shows a considerably smaller number of proliferative cells than the pallium. Despite we cannot conclude the existence of separate neurogenic niches, we have observed differences in the density of PCNA-ir cells in the VZ of different pallial regions, with higher proliferation in the ventral pallium. Interestingly, in zebrafish, the pallium presents a smaller number of proliferating cells than the subpallium, where cells are densely grouped (Grandel et al. 2006; Mueller and Wullimann, 2016). It has been proposed that structures that become postmitotic late in development become more enlarged that structures that are born earlier, that is, that the telencephalon in vertebrates tends to be disproportionally large because neurogenesis in this region is generally protracted (see Striedter and Charvet, 2008 and references therein). Early *vs*. delayed and extended telencephalic neurogenesis in birds has been related to precociality *vs*. altriciality (Charvet and Striedter, 2011). The altricial-precocial spectrum describes the degree of morphological maturation of offspring at the moment of hatching (Álvarez-Hernán et al. 2019). In fish, precocial species hatch at an advanced stage of development, while altricial species hatch at a less developed stage. Though *S. canicula* is considered a precocial fish, delayed neurogenesis in the adult telencephalon to produce GABAergic neurons that could migrate and become integrated in the olfactory bulbs could be related to the disproportionate growth of this structure relative to brain with respect to that observed in other vertebrates (Yopak, 2012). However, it remains unexplored if this difference in the proliferation pattern in different telencephalic (pallial vs subpallial) domains results in different allometric relationships of these domains to overall telencephalon size. Besides control of neurogenesis timing, other mechanisms have been proposed to be involved in adult brain size including changes in regionalization during early development and apoptosis within proliferative zones (see Charvet and Striedter, 2008; Striedter and Charvet, 2008).

In addition, as in other fish (Zupanc et al. 2006), we have found many scattered cells throughout the telencephalon of *S. canicula* in adventricular positions, which suggest us that neuroblasts maintain their progenitor capacity time after they leave the VZ. The fact that sharks exhibit high levels of proliferation in the telencephalon in juveniles and adults (present results) supports the idea that the number of neurogenic niches and proliferating cells increases in anamniotes (Ganz and Brand, 2016). However, in contrast to what it could be expected, in lampreys (an ancient, extant lineage of jawless fish) PCNA immunoreactivity disappears at the end of the developmental period (Villar-Cheda et al. 2006). Despite in larvae the number of proliferating cells is significant, after metamorphosis and in the adulthood no proliferating cells have been detected under normal conditions. The presence of a high number of proliferating cells in the telencephalon of the catshark and the absence of proliferation in adult lamprey point to an evolutionary origin of adult neurogenesis in vertebrates in the transition from agnathans to jawed vertebrates.

### aNSCs in the telencephalon of juvenile sharks’ express radial glia markers and show proliferative activity

In the adult neurogenic niches of mammals, many cellular types coexist. One of these cells are radial glial-like cells (B cells in the SVZ and Type-1 in the SGZ). These cells have a glial nature and act as progenitor cells in the first steps of the neurogenic process, though they are relatively quiescent (see Introduction and Fig. 7). They express glial markers such as GFAP, BLBP, GS, and stem cell markers such as Sox2, among others (Doetsch et al. 1997, 1999; Seri et al. 2004; Ming and Song, 2011; Götz, 2013; Bond et al. 2015; Bonaguidi et al. 2016; Lim and Alvarez-Buylla, 2016). In birds, studies using electron microscopy, incorporation and examination of tritiated thymidine (3HT; a mitosis marker) labeled cells have allowed identification of radial glial cells in the VZ of the telencephalon with stem cell and neurogenic capacity (Álvarez-Buylla et al. 1990, 1998). In reptiles, similar electron microscopy studies combined with BrdU and 3HT have identified proliferating radial glial cells in the VZ of the telencephalon of lizards (García-Verdugo et al. 1981; Pérez-Cañellas and García-Verdugo, 1996; Font et al. 2001; Grandel and Brand, 2013). Besides, studies in turtles also show radial glial cells in the VZ of the adult telencephalon (Clinton et al. 2014). However, these studies do not mention the proliferation rate of these cells. In amphibians, studies in newts have evidenced that the entire VZ is formed by radial glial cells containing GFAP or GS that additionally coexpressed Sox2. These cells also express proliferating markers in the “hot spots”, and they have neurogenic potential (Kirkham et al. 2014). However, outside the “hot spots”, radial glial cells mainly remain in a quiescent state.

In teleost fish, a detailed study in the telencephalon of zebrafish has identified numerous radial glial cells that express glial markers such as BLBP or GFAP, and Sox2 (a marker of stem cells; März et al. 2010). Some of these cells are PCNA-ir and incorporate BrdU at slow rates, which means that they have proliferative capacity. However, most Sox2-expressing radial glial cells are PCNA-negative, which have been interpreted as quiescent progenitors, a feature of mammalian B cells.

In the VZ of the telencephalon of the catshark, most of the ventricular cells showed a radial morphology and contain GFAP, BLBP and GS. As in zebrafish, we have observed numerous cells that express *ScSox2* and are PCNA-positive. In addition, we also found *ScSox2* expressing cells that do not show proliferative activity, which probably represent quiescent progenitor cells. Our results are quite similar to that reported in teleost where quiescent and proliferating subtypes of progenitor radial glial cells have been described (März et al. 2010).

Lampreys also exhibit radial glial-like cells in the adult telencephalon. These radial cells extend processes form the VZ to the pia (as in amphibians and fishes), but they express cytokeratins instead of glial markers such as GFAP (Merrick et al. 1995). However, no PCNA positive cells have been found in the telencephalic VZ of this species (Villar-Cheda et al. 2006), suggesting that adult radial glial-like cells in lampreys do not have proliferative potential. Of note, studies about proliferation in larvae of lampreys have shown that seasonal changes or lesions lead to a reactivation of proliferation in cytokeratin containing cells in the rombencephalon or spinal cord (Zhang et al. 2014), supporting an ancient origin of radial glial-like cells as progenitor cells in the postnatal neurogenic process.

### Heterogeneity of radial glial cells

An heterogeneous proliferation rate of adult pallial radial glia has been reported in zebrafish (März et al. 2010). Whether this heterogeneousness reflects the existence of progenitor radial glia subtypes, a hierarchy in their recruitment cascade or stochastic variations within a single cell population has not been determined (März et al. 2010). Here we have used the radial glial markers GFAP, BLBP and GS to evaluate if different subpopulations of radial glial cells present in the VZ are associated to different proliferation rates. Recently, GFAP, BLBP or GS positive cells with morphology of radial glia have been previously reported in this species (Docampo-Seara et al. 2019 and references therein), but its proliferative potential has never been investigated in juvenile specimens. Using double immunofluorescence in the catshark we have seen that BLBP and GS label the same population of cells. Curiously, double immunofluorescence between GFAP and GS has shown some positive isolated cells for GFAP or GS (very rare). Due to the scarcity of these cells and their position of one respect to the other, we think that they may correspond to newborn radial glial cells that are generated in order to expand the ventricular zone with the natural growth of the individual, and they are not a subtype of radial glial cells. Due the fact that PCNA immunoreactivity was found in subgroups of both GS/BLBP-ir cells and GS/BLBP/GFAP-ir cells we cannot associate a quiescent *vs*. a proliferative state with the expression of any of the glial markers used in this study.

Recent studies have shown that the VZ of juveniles of catshark present a considerable number of cells expressing GFAP, BLBP and GS (Docampo-Seara et al. 2019). Surprisingly, in the present study we have found that these cells, in addition, coexpress S*cSox2* (presumptive progenitor radial glia). This fact may be interpreted in two different ways: (1) either radial ependymoglia express *ScSox2* (i.e. differentiated radial glial cells retain neurogenic potential as has been observed in Müller glial cells in the retina of sharks; unpublished data); (2) all *ScSox2* expressing cells in the telencephalon of juveniles are radial glia progenitor cells, and differentiated radial ependymoglia will differentiate latter in adulthood. Since in sharks, as in zebrafish (Than-Trong and Bally-Cuif, 2015), GFAP, BLBP and GS do not allow to differentiate radial glia progenitor cells from radial ependymoglia, future investigations must be directed towards the search of specific molecular markers of these cell types in this species.

### Intermediate progenitor cells based on non-radial morphology, transit-amplifying features and/or neuronal commitment

Once radial glial cells reactivate their cellular cycle and undergo mitosis, they generate fast cycling progenitors that subsequently generate neuroblasts. Fast cycling progenitors are also called IPCs, transit amplifying cells or non-radial progenitors (Fig. 7). In mammals these IPCs are located close to the radial glial progenitors and do not leave the neurogenic niche. IPCs have not been found in the VZ of the lateral ventricles of the brain of adult birds by means of electron microscopy and 3HT incorporation assays (Álvarez-Buylla et al. 1998; reviewed in García-Verdugo et al. 2002). In reptiles, the existence of adult IPCs has not been proved. On the contrary, in amphibians, drug treatment with AraC (an IPC killer) and BrdU incorporations have clearly evidenced the existence of cells with transit-amplifying features (Kirkham et al. 2014). In teleost fish, experiments with BrdU incorporations have evidenced the existence of fast dividing cells, that decrease their expression of glial markers and start to express neuroblast markers such as PSA-NCAM, which match the definition of IPCs (März et al. 2010), though the existence of this cell type in zebrafish is underrepresented.

In mammals, these fast cycling progenitors can be additionally identified by their decreased expression of glial markers and increased expression of neuronal commitment markers such as Tbr2 (glutamatergic lineage marker) or Dlx2 (GABAergic lineage marker) (Doetsch et al. 1997, Ming and Song, 2011; Hodge et al. 2012; Bond et al. 2015; Lim and Alvarez-Buylla, 2016). During telencephalic development, glutamatergic neurons are born locally in the pallial VZ from radial glial progenitors from early stages (around E11) until around E17, when pallial-born neurogenesis decays to barely undetectable levels (Englund et al. 2005; Arnold et al. 2008). GABAergic interneurons are produced in the subpallial VZ, also form radial glial progenitors, following a similar timing (between E11 and E17; Sultán et al. 2013) and migrate tangentially to the developing pallium (first described by Anderson et al. 1997). A similar separation in the origin of the glutamatergic and GABAergic cell lineages has been reported during early telencephalic development in *S. canicula* though (Quintana-Urzaiqnui et al. 2015; Docampo-Seara et al. 2018).

In mammals, adult B cells inherit the regional signature of embryonic radial glial progenitors, therefore the adult SGZ (pallium) mainly produce glutamatergic projection neurons and progenitors in the adult SVZ (subpallium) produce GABAergic neurons (Fuentealba et al. 2015). However, it has been shown that progenitors within the same location at different developmental times (embryonic *vs*. adult) slightly differ in the types of neurons they generate (Obernier and Álvarez-Buylla, 2019). Thus, the adult SVZ can produce a subset of glutamatergic neurons that migrate through the RMS to the olfactory bulbs (Hodge et al., 2012) while the case of the pallium can generate a specific subset of GABAergic interneurons that migrate to the olfactory bulb (Kohwi et al. 2007; Merkle et al. 2007; Ventura and Goldman, 2007; Cai et al. 2013).

In the present work we have investigated the existence of IPCs in the adult ventricular zone of *S. canicula* by using double IHQ against PCNA and ISH for *ScTbr2* (glutamatergic IPCs) and *ScDlx2* (GABAergic IPCs). We did not find *ScTbr2* expression in the VZ of the telencephalic ventricles, but rather in the intermediate zone, being especially abundant in the ventral pallium. This contrasts with results in adult zebrafish, where Tbr2 and Prox1 positive cells (both markers of glutamatergic cells) have been reported in different domains of the pallial VZ (Ganz et al. 2015). The absence of *ScTbr2* in the VZ of the catshark (present results) suggests that either IPCs (which are located in/close the ventricle) are not present in the pallium or they do not express *ScTbr2*, which evidences the need of specific markers for glutamatergic IPCs. Further studies using cell-tracking are needed to clarify this point. In contrast we found *ScDlx2* positive cells ventricularly, many of them also positive for PCNA, suggesting the presence of IPCs of the GABAergic lineage in the pallial VZ. Interestingly, BrdU birthdating assays showed that some actively proliferating cells (that incorporate BrdU) were also GABA immunoreactive. Whether these dorsal pallial progenitors generate a specific subset of GABAergic interneurons migrating to the olfactory bulb, as in mammals, deserves further investigation.

### Migratory neuroblasts

The third progenitor cell type in mammals is a population of migratory neuroblasts (Fig. 7) that, in addition to Tbr2 or Dlx2, express DCX or PSA-NCAM and can exit the neurogenic niche to reach their final destination in the telencephalon (Doetsch et al. 1997). In birds, the presence of IPCs has not been demonstrated but the increase of DCX immunoreactivity in the adult brain suggest the existence of migratory neuroblasts (type A) derived from radial glial (type B) progenitors (Barnea and Pravasudov, 2011; Mazengenya et al. 2018). In reptiles, DCX positive cells have been detected in the adult brain of crocodiles (Ngwenya et al. 2017) and turtles (Macedo-Lima et al. 2016). In amphibians, the high levels of PSA-NCAM surrounding the proliferative ventricle clearly support the presence of neuroblasts (Kirkham et al. 2014). In zebrafish PSA-NCAM positive cells have been found close to the VZ, which have typical morphology and markers of neuroblasts (März et al. 2010). In this study we found both *ScDlx2* and *ScTbr2* out of the VZ. Besides, DCX positive cells in the ventricular and in subventricular positions, some of them also immunoreactive to PCNA. This clearly suggests that proliferating cells in the mature brain of sharks give rise to new neurons.

### The neurogenic niche of sharks and mammals: continuous neurogenesis vs. reactivation of silenced progenitors

During early development the neural ectoderm is comprised of neuroepithelial cells (NECs) that undertake self-renewing symmetric divisions that increase the size of the precursor cell pool (Fig. 8A). Later in development, NECs begin to express glial molecular markers and become radial glial cells. These cells are located in the VZ where they initially undergo symmetric divisions that produce additional radial glial cells and expand the proliferative population in the VZ. Later, these cells undergo asymmetric divisions to generate one self-renewed radial glial cell and one daughter neuron (reviewed in Götz and Huttner, 2005). In the cortex of mammals, they can also give rise to neurons or oligodendrocytes indirectly by generating progenitors that migrate to the SVZ, where they divide symmetrically to produce two daughter cells. Progenitors that divide in the SVZ are known as basal or intermediate progenitors. Following their final division at the VZ, radial glial cells become detached from the ventricle and subsequently are translocated towards the pial surface, where they differentiate into astrocytes (reviewed in Martínez-Cerdeño and Noctor, 2018).

**Figure 8.**
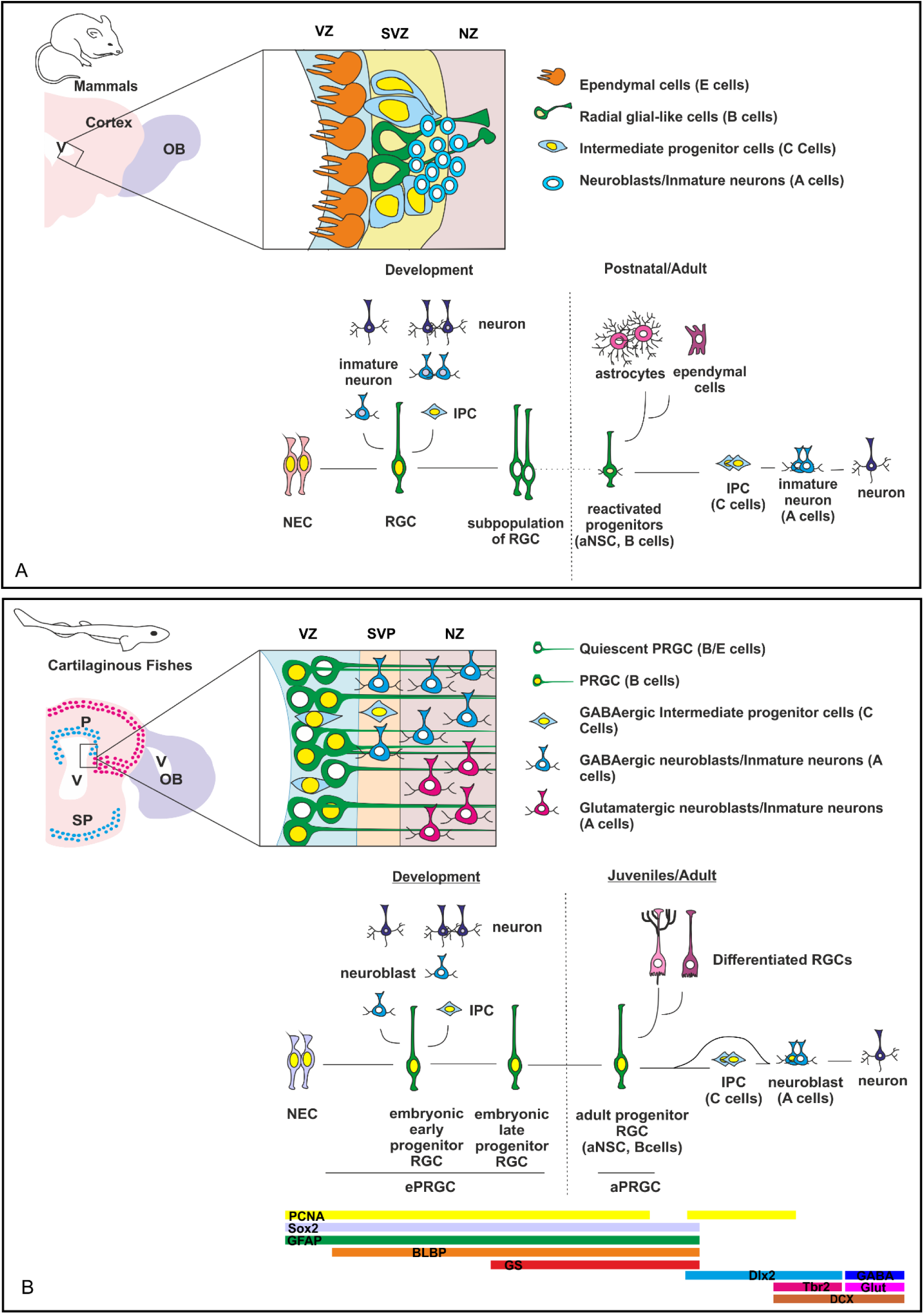
Schema showing the cell organization of the neurogenic niche of adults and the neurogenic process from early development to adults in mammals **(A)** and sharks **(B).** The schema of the mammalian neurogenic niche was based on data from Ming and Song, (2011); the information/design of the progression from embryo to adult was based on data from Kriegstein and Álvarez-Buylla, (2009). Abbreviations: aPRGC: adult progenitor radial glial cell; aNSC: adult neural stem cell; ePRGC: embryonic progenitor radial glial cell; IPC: intermediate progenitor cell; NEC: neuroepithelial cell; NZ: neurogenic zone; OB: olfactory bulb; P, pallium; RGC: radial glial cell; SP, subpallium; SVP, subventricular position; SVZ: subventricular zone; V: ventricle; VZ: ventricular zone.

In mammals, radial glial cells are lost in most brain regions at the end of the neurogenic period, either by symmetric self-consuming neurogenic divisions or by becoming glial cells (Paridaen and Huttner, 2014). However, reactivated progenitor cells, both in the SVZ and in the SGZ, exhibit morphologies and molecular markers typically expressed by radial glial cells and astrocytes that lead to suspect that progenitor cells in the adult brain may correspond to embryonic radial glial progenitors that persist in adulthood (see Fuentealba et al. 2015). Indeed, it has been shown that these cells are generated from a subpopulation of progenitor radial glial cells in the embryo. These cells remain quiescent until they become reactivated at different ages in the postnatal brain, leading to a neurogenic process that recapitulates the embryonic neurogenesis (Kriegstein and Alvarez-Buylla, 2009; Berg et al. 2019; Obernier and Álvarez-Buylla, 2019; Fig. 8A). These subpopulations of cells only persist in particular regions of the telencephalon and therefore neurogenic niches in mammals correspond to well defined areas.

This situation differs from what is observed in sharks (Fig. 8B), since abundant progenitor cells are observed through the VZ of the pallium during the whole life, resembling the situation described in teleost fish (reviewed in Than-Trong and Bally-Cuif, 2015). Whether this proliferation pattern is the result of continuous neurogenesis or correspond to reactivation of silenced progenitors has not been explored. However, cell types (identified by morphology, position or neural commitment markers) observed in the embryo are similar to those found in the adult. In fact, we have previously shown that the VZ of the telencephalon is highly proliferative even at late stages of embryonic development (i.e., PCNA and PH3 positive cells have been described along the entire VZ of the telencephalon; Quintana-Urzainqui et al. 2015). In addition, a deep characterization of these cells has shown that most proliferating cells present in the ventricular zone correspond to radial progenitor cells that expressed molecular glial markers as GFAP, BLBP and GS (Docampo-Seara et al. 2019). Results obtained in embryos are highly coincident with those obtained in juvenile/adults (present work), evidencing that the VZ of the posthatching catshark show a cell organization similar to that observed in embryos. This fact indicates that in contrast with mammals, radial glial cells with embryonic molecular features persist in the telencephalic VZ of the catshark, leading to a telencephalic neurogenic niche which is not restricted to a particular ventricular region, but to the entire telencephalic VZ (Fig. 8B).

## Conclusions

We have investigated adult neurogenesis under an evolutive perspective by characterizing the neurogenic niche in the telencephalon of catshark. The phylogenetic position of cartilaginous make this group essential in comparative studies to infer the ancestral condition of vertebrate neurogenesis. High rates of proliferation were found in the telencephalic VZ of the catshark, in contrast to lampreys, which points to an evolutionary origin of adult neurogenesis in vertebrates in the transition from agnathans to gnathostomes. We also have shown that the VZ exhibits high numbers of progenitor cells that matches the definition of a B cell (radial and mainly quiescent cells that express *ScSox2* and radial glial cell markers). Besides, we have pointed to the existence of putative C cells (IPCs of GABAergic nature). Finally, some DCX-immunoreactive neuroblasts were found to be proliferative (immunoreactive to PCNA), which matched the definition of A cells. Therefore, we show that the main types of cells found in the mammalian telencephalic niche are already present in cartilaginous fish. This study constitutes an important step in the race to unravel the evolution of adult neurogenesis in vertebrates.

## Acknowledgments

Authors declare not to have conflicts of interest. We would like to thank the Aquarium of “O Grove” for kindly providing juveniles samples of catsharks. We would also like to thank Prof. Dr. Elisabeth Pellegrini for her advice during the elaboration of this manuscript. This work was supported by the Spanish Ministerio de Economía y Competitividad-FEDER (BFU2014-5863-1P) and Ministerio de Ciencia e Innovación-FEDER (BFU-2017-8986-1P)

## Disclosure of potential conflicts of interest

### Ethical approval

All procedures conformed to the guidelines established by the European Communities Council Directive of 22 September 2010 (2010/63/UE) and by Spanish Royal Decree 1386/2018 for animal experimentation and were approved by the Ethics Committee of the University of Santiago de Compostela.

## Supplementary Figure legends

**Supplementary Figure 1.** Schema showing the anatomy of the telencephalon of the catshark, showing topographic (A-C) and topologic (D-E) views. In the schemas from telencephalic transverse sections, pallium is represented in red and subpallium is represented in green. Schemas based on Smeets et al. 1983 and Santos-Durán et al. 2015.

**Supplementary Figure 2.** Schema from transverse sections of the telencephalon of juveniles (Juv) and adults representing the box area used for cell counting of PCNA-ir cells in the ventral pallium, the number of samples used and the average and standard deviations. Numbers of cells were quantified in the ventral pallium, where high numbers of PCNA-ir cells are observed in juveniles with respect to other telencephalic regions.

**Supplementary Figure 3.** Schemes and photomicrographs after BrdU 24h pulses in the telencephalon of catshark juveniles (Juv) showing double labelled cells for BrdU and GAD in the VZ of the dorsal and medial pallium **(A-A’’)** and labelled cells for BrdU but not for GAD in the subpallial ventricular zone **(B-B’’). N**ote the high expression of GAD in the subpallium at the level of the basal superficial area **(B-B’’).** Scale bars: 50 µm (A-A’’); 200 µm (B-B’’). Abbreviations: BSA, basal superficial area; DP, dorsal pallium; MP, medial pallium; SP, subpallium; V, ventricle.

