## Supplementary figures and images for "Characterization of neurogenic niches in the telencephalon of juvenile and adult sharks"

### Supplementary Figure 1

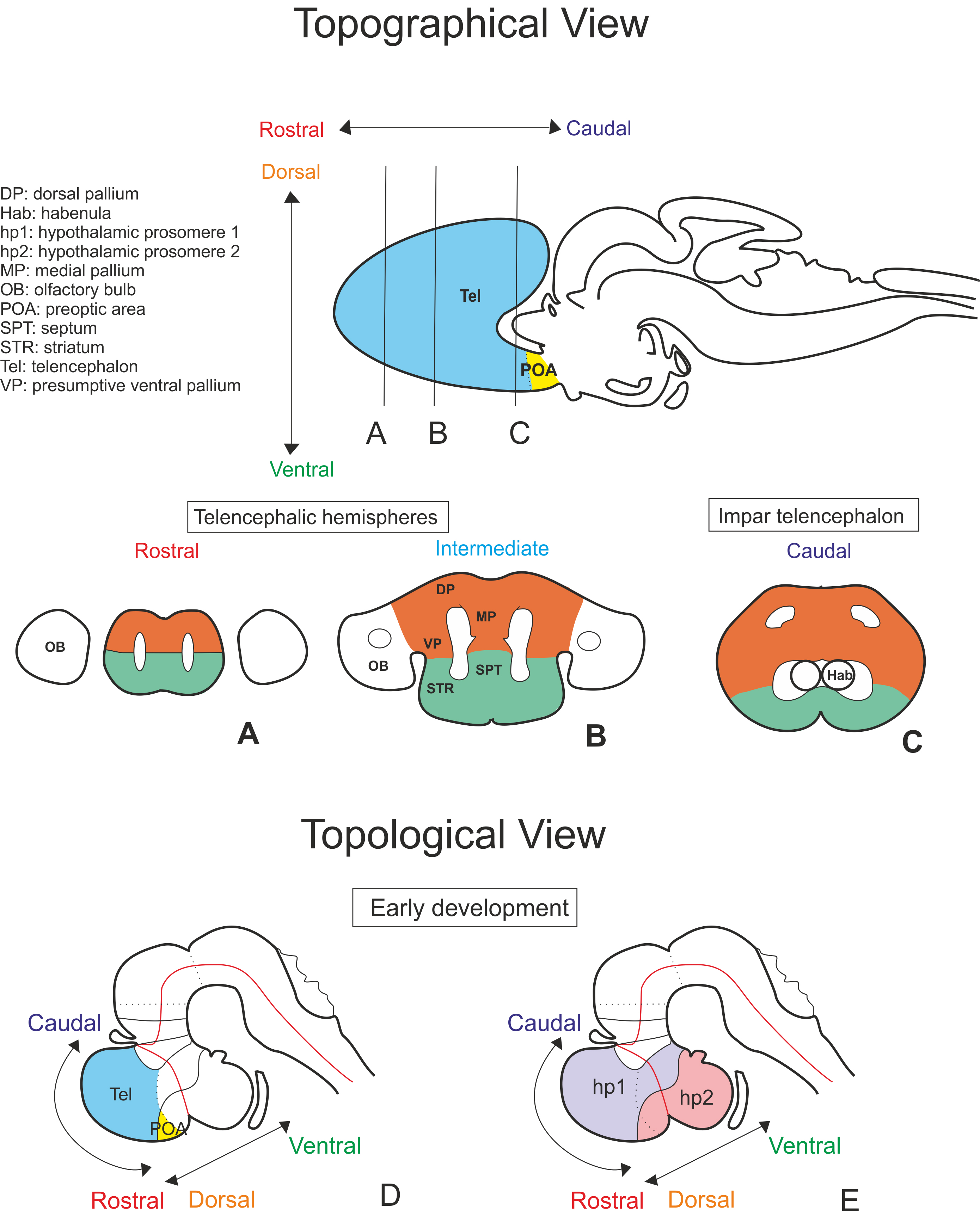

### Supplementary Figure 2

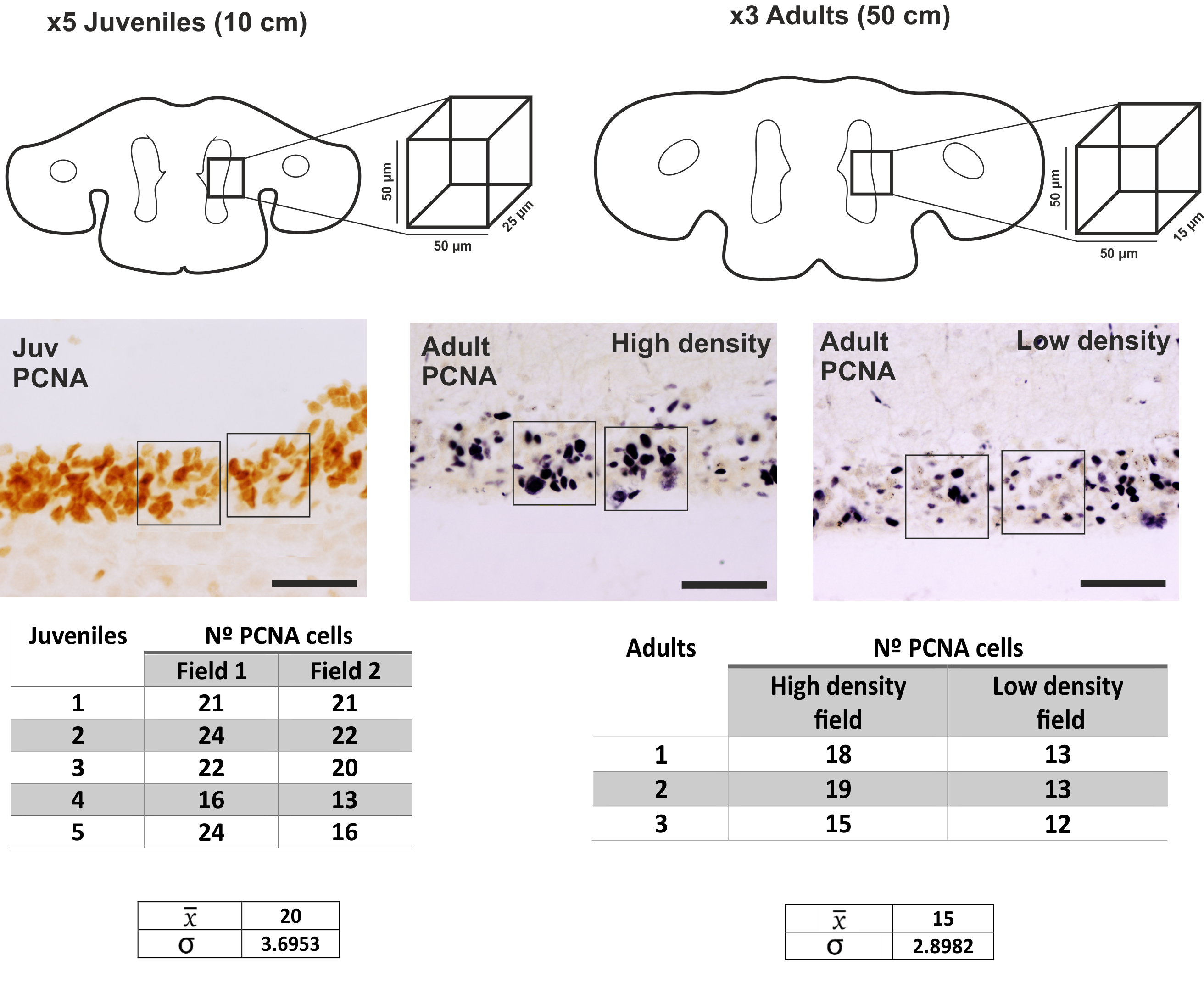

### Supplementary Figure 3

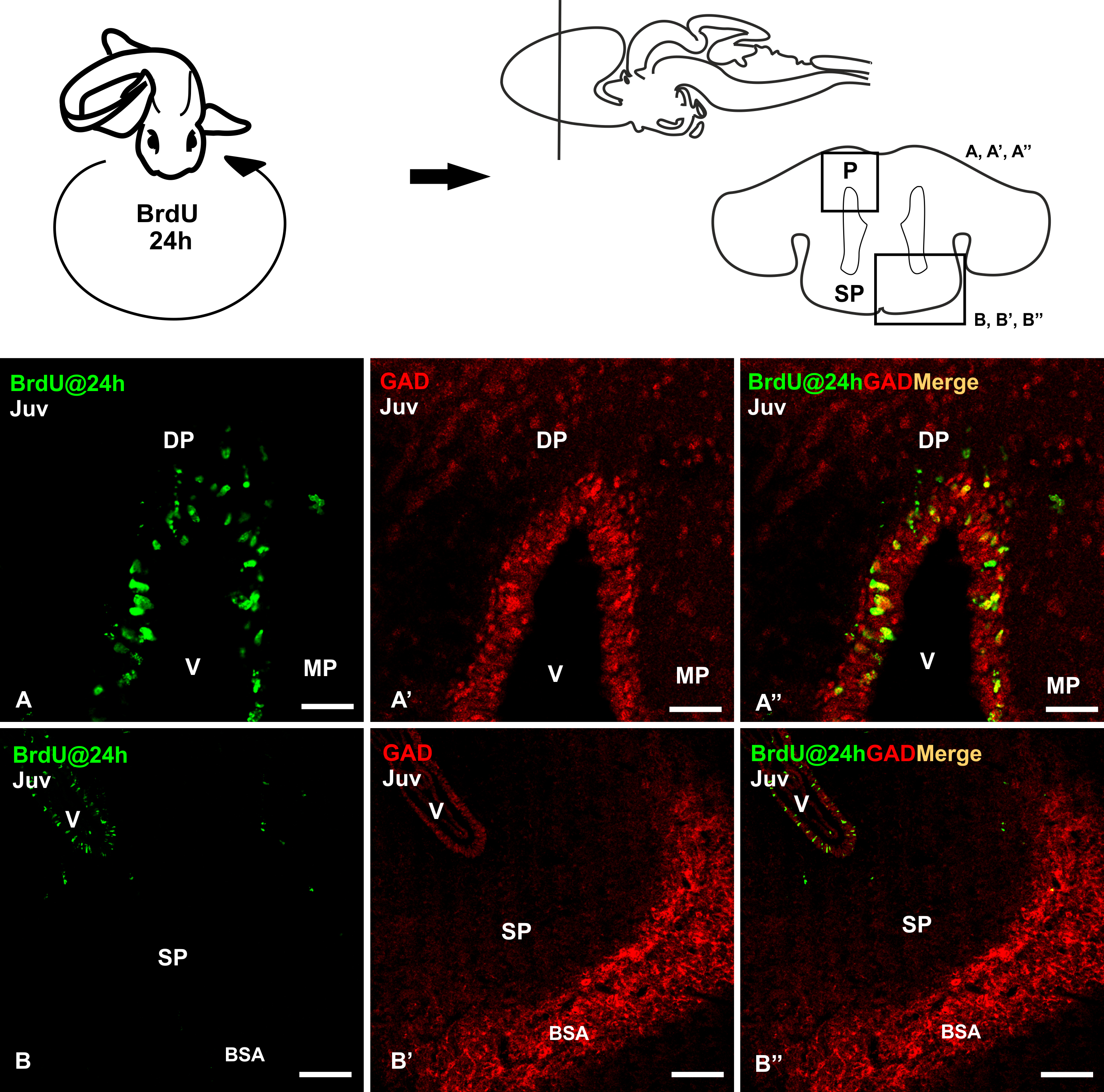
